# ddRAD-seq-derived SNPs reveal novel association signatures for fruit-related traits in peach

**DOI:** 10.1101/2023.07.31.551252

**Authors:** Najla Ksouri, Gerardo Sánchez, Carolina Font i Forcada, Bruno Contreras-Moreira, Yolanda Gogorcena

**Author notes:** Senior authors (Bruno Contreras-Moreira, Yolanda Gogorcena). Emails: Najla Ksouri, Gerardo Sánchez, Carolina Font i Forcada, Bruno Contreras-Moreira*, Yolanda Gogorcena.

## Abstract

Breeding for new peach cultivars with enhanced traits is a prime target in breeding programs. In this study, we used a discovery panel of 90 peach accessions in order to dissect the genetic architecture of 16 fruit-related traits. ddRAD-seq genotyping and the intersection between three variant callers yielded 13,045 high-confidence SNPs. These markers were subjected to an exhaustive association analysis by testing up to seven GWAS models. Blink was selected as the most adjusted, simultaneously balancing false positive and negative associations. Totally, we identified 16 association signals for six traits showing high broad-sense heritability: harvest date, fruit weight, flesh firmness, contents of flavonoids, anthocyanins and sorbitol. By assessing the allelic effect of significant markers on phenotypic attributes, nine SNP alleles were denoted favorable. A promising marker (SNC_034014.1_7012470) was found to be simultaneously associated with harvest date and fruit firmness conferring a positive allelic effect on both traits. We anticipate that this marker could be used to improve firmness in late harvested cultivars. Candidate causal genes were shortlisted when fulfilling the following criteria: i) position within the linkage disequilibrium block, ii) functional annotation and iii) expression pattern. A bibliographic review of previously reported QTLs mapping nearby the associated markers allowed us to benchmark the accuracy of our approach. Despite the moderate germplasm size, ddRAD-seq allowed us to produce an accurate representation of peach’s genome resulting in SNP markers suitable for empirical association studies. Together with candidate genes, they lay the foundation for further genetic dissection of peach key traits.

## Background

Peach is one of the most economically valued fleshy fruits worldwide (FAO, http://faostat.fao.org). The advances in the peach industry largely rely on fruit quality improvement in response to the market and consumers’ demands. The term quality may include all agronomical aspects and chemical compounds such as fruit size, firmness, sugar and acid concentration, etc. Some of those characteristics are thought to be monogenic, controlled by a single gene (fruit shape, hairiness, flesh color, texture)^1–3^ while others are polygenic, such as sugar content, fruit firmness, antioxidant concentration^4^.

Breeding for polygenic quantitative traits is far from being a straightforward task. Thus, insights on genetic drivers controlling these traits and their inheritance are required to bridge the phenotype-genotype gap^3, 5^. For instance, the development of molecular markers linked to desirable traits would considerably speed up the selection of superior plant varieties through marker-assisted selection (MAS)^6^. Genome-wide association studies (GWAS) have also revolutionized the breeding process by detecting the genetic loci underlying trait variations at a relatively high resolution. This approach has been successfully applied in many breeding programs. For instance, GWAS have provided insight into fruit-related traits such as skin color in apple^7^ and fruit firmness in sweet cherry^8^. The power and prediction accuracy of GWAS critically depend on various considerations, including phenotypic data quality, experimental sample size, linkage disequilibrium (LD) between genetic variants and population structure. If not adjusted properly, these factors may lead to spurious associations as well as masking the true ones. Another key factor while performing GWAS is the density and chromosome distribution of markers/SNPs along the reference genome.

Generally, genotyping methods fall into three categories; whole genome resequencing, reduced representation sequencing, and SNP arraysLL^9^. Whole genome resequencing returns the highest number of SNP calls if sequencing depth is sufficient, which is expensive for large genomes. For this reason, SNP arrays are widely used, reducing the cost and enabling the detection of thousands of SNPs in a single assay^9^. In peach, commercially available arrays IPSC peach 9K^10^ and IPSC peach 18K^11^ have been used to explore the genetic diversity and to assist the breeding process^1, 12^. Despite their utility, the major drawback of SNP genotyping arrays consists in their ascertainment bias^13^. In other words, they narrow the discovery of novel variants other than those detected in the discovery panel and used to build the respective array. This might distort subsequent genetic inferences. Additionally, efficient SNP probes require a well-assembled reference genome and their design and further optimization can be time consuming.

With the massive progress of high-throughput technologies, reduced representation sequencing such as restriction-associated DNA (RAD) sequencing and its derivative (ddRADseq) emerged to overcome both cost and ascertainment bias^14^. Double digest restriction-site associated DNA (ddRADseq) relies on the use of a pair of restriction enzymes to limit the sequencing effort to a subset of evenly distributed loci in the genome^14^. Moreover, by picking the best enzyme combination, repetitive DNA can be less targeted, thereby reducing the computational burden associated with aligning genomes with highly repetitive segments.

Unlike other genotyping methods, prior genomic information is strictly not required for ddRADseq^14^. Nevertheless, as shown in this work, it is most powerful when combined with a reference genome sequence. From a technical standpoint, a common shortcoming of ddRADseq is the high rate of missing calls which can be straightforwardly handled through genotype imputation.

Herein, we report the application of ddRADseq genotyping to identify high confidence SNPs in a discovery panel of 90 *Prunus persica* accessions. Consequently, GWAS was carried out to identify genomic loci associated with 16 fruit traits. To optimize the analysis and to overcome the limitations arising from the size of our peach germplasm, we considered the following aspects: 1) peach accessions were geographically distant in order to maximize the genetic variance, 2) SNPs were called using three variant detectors (BCFtools, Freebayes and GATK) and only those resulting from the intersection were retained for subsequent analysis, and 3) several statistical models were assessed to control the confounding effects.

Genotype-to-phenotype associations for agronomic and fruit-related traits have been widely tested in peach using different genotyping methods like SSRs^15^, 9K SNP array^1, 4, 16^, 18K SNP array^3, 12^ and high-throughput resequencing technology^17^. However, to the best of our knowledge this is the first report characterizing the genetic architecture of peach traits using ddRADseq-derived SNPs. In this study, we propose best practices for GWAS analysis mainly relying on a comparative approach for SNPs calling and statistical model assessment. Therefore, we demonstrate the utility of ddRAD-based genotyping in unveiling desirable alleles and genomic regions putatively responsible for trait variation. By contrasting our findings with those previously reported using the peach 9K SNP array^16^ we confirm the accuracy of our approach.

## Results

### Phenotypic analysis and heritability

Broad sense heritability was estimated over three consecutive years and the results denote that most of the traits were highly heritable (**Figure 1.A**). Hence, their phenotypic variability among the individuals was mainly driven by the genetic effects. However, contents of glucose, fructose, sucrose and total sugars (TS) were found to be lowly heritable traits (H^2^ < 0.5), denoting that their variability may be mostly due to the environmental factors. These traits were therefore left out of the association analyses. Furthermore, normal distribution fit tests conducted on averaged phenotypic measures, revealed that six out of 16 traits were found to be normally distributed (flesh firmness, soluble solids content (SSC), ripening index, vitamin C, relative antioxidant capacity (RAC) and glucose). Source code, documentation and detailed results can be accessed at https://github.com/najlaksouri/GWAS-Workflow. The remaining ones, skewed either positively or negatively, were transformed accordingly. Likewise, the phenotypic correlation was estimated and significant interactions between agronomical and fruit quality traits were observed (**Figure 1.B**). For instance, harvest date (HvD) had the highest heritability estimates (H^2^=0.94) and exhibited strong positive correlations with flesh firmness, sugar contents measured as (SSC, TS and sorbitol) and antioxidant activity measured as (RAC, flavonoids and phenols). As expected, moderate positive interaction was also reported between the HvD and fruit weight as well as between total and individual sugars. Moreover, a strong positive correlation was also observed between total phenolics and flavonoids. Indeed, flavonoids are the largest group of naturally occurring phenolic compounds in plants. Both compounds showed a significant positive interaction with (RAC) suggesting that they could be used as a good indicator of antioxidant properties in peaches.

**Figure 1.**
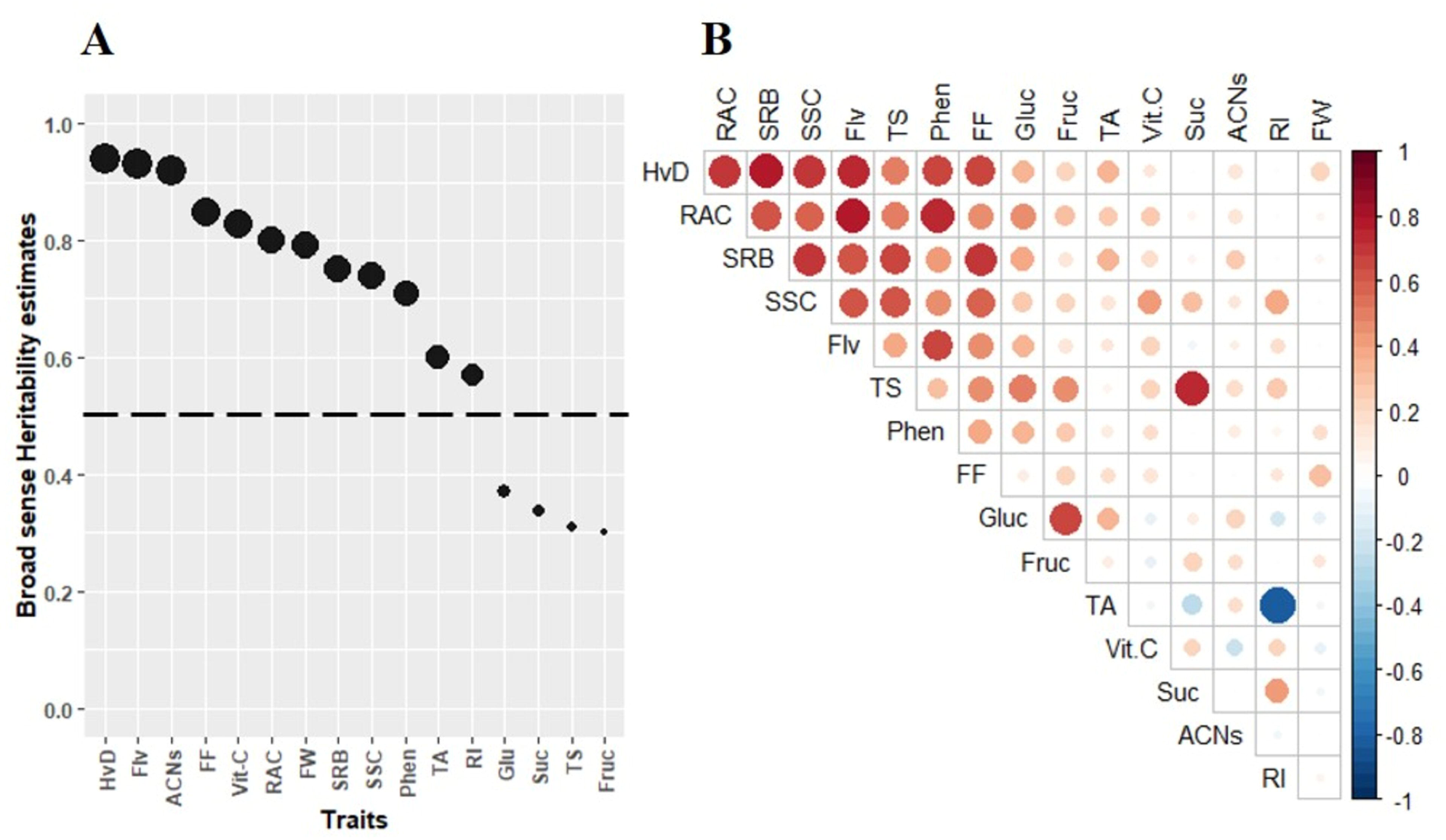
**(A):** Broad sense heritability estimates and (**B**): phenotypic correlation among 16 peach agronomical and fruit quality traits. Dashed horizontal line corresponds to heritability threshold (H^2^ = 0.5). Correlation between traits was estimated using *Pearson* correlation. Significant positive and negative correlations are displayed in red and blue respectively (*P* < 0.05). Color intensity and size of the circles are proportional to the correlation coefficients. Abbreviations are as follows: harvest date (HvD), fruit weight (FW), flesh firmness (FF), soluble solids content (SSC), titratable acidity (TA), ripening index (RI), content of vitamin C (Vit C), total phenolics (Phen), anthocyanins (ACNs), sucrose (Suc), glucose (Glu), fructose (Fruc), sorbitol (SRB) total sugars (TS) and relative antioxidant capacity (RAC).

### SNP genotyping

To construct an informative SNP panel, polymorphic sites were called in individual sample mode using three different algorithms. Raw calls were subjected to standardized quality thresholds in order to mitigate the effect of sequencing and/or alignment flaws. Post-filtered calls from each pipeline were merged together into multi-samples format (**Table 1**). According to our results, GATK-HaplotypeCaller (HC) outperformed both Freebayes and BCFTools in terms of computational time and sensitivity yielding a total of 233,535 SNP calls (see repository https://github.com/najlaksouri/GWAS-Workflow). Freebayes ranked second, followed by BCFTools, with 166,080 and 148,998 SNPs, respectively. For a robust variant detection, the intersection between multi-sample sets was computed. About 32% of SNPs were found to be commonly shared by the above-stated tools. Multi-allelic and scaffold variants were excluded and additional filters (missing call rate and MAF) were applied (**Table 1**). Finally, a set of 13,045 SNPs was kept for subsequent analysis.

**Table 1.** SNPs count and filtering steps.

| Applied filters | Retained SNPs |
| --- | --- |
| Clean multi-samples SNPs from GATK-HaplotypeCaller | 233,535 |
| Clean multi-samples SNPs from Freebayes | 166,080 |
| Clean multi-samples SNPs from BCFtools | 148,998 |
| Intersected set | 56,430 |
| Removing scaffold SNPs | 56,647 |
| Removing multi-allelic sites | 56,430 |
| Missing call rate < 20% | 26,188 |
| Minor Allele Frequency > 0.05 | 13,045 |

Using VEP tool, polymorphic sites were found to be distributed along upstream (21%), downstream (9%), intronic (26%) and intergenic (8%) regions (**Figure S1**). Low proportions of SNPs were tagged as 3’ UTR and 5’ UTR variants. Within coding regions, 11% of SNPs were defined as synonymous while 13% were annotated as missense variants.

### SNP distribution and LD decay

The distribution of polymorphic sites was calculated within adjacent windows of 1 Mbp and provided a genome-wide coverage estimate along the eight peach chromosomes. As illustrated in **Figure 2.A**, markers were unevenly partitioned throughout the genome with the highest number of mapped SNPs on chromosome 2 (4,440) and the lowest on chromosome 5 (1,768). Interestingly, SNPs accumulated within the short arms of chromosomes 2 and 4. In contrast, large gaps were observed towards the telomere of the long arm of chromosome 2. Similarly, several blank regions were located along chromosome 1. Gaps highlighted with asterisks correspond to predicted centromeric regions^18^.

**Figure 2.**
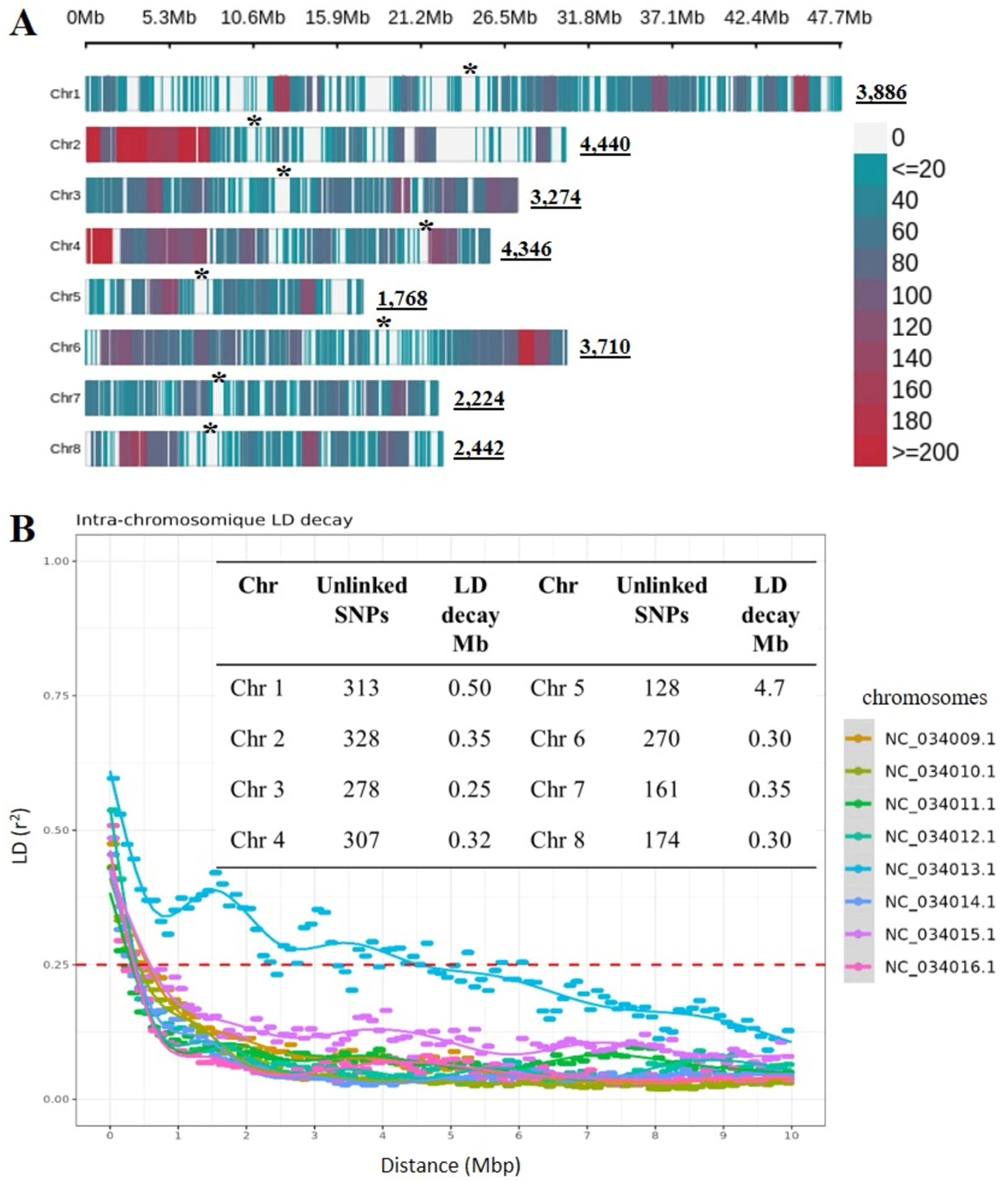
SNPs density plot and intra-chromosomique linkage disequilibrium decay. (**A**): SNPs density across the eight peach chromosomes. The horizontal axis shows the chromosome length in (Mbp) and the different colors reveal the SNP density per window of 1 Mbp. Underlined numbers correspond to the total number of polymorphic sites per chromosome. The asterisks highlight the putative position of centromeres predicted as follows: Chr 1=NC_034009.1: (∼21 Mbp), Chr 2=NC_034010.1: (∼8 Mbp), Chr 3 =NC_034011.1: (∼12 Mbp), Chr 4=NC_034012.1: (∼24 Mbp), Chr 5=NC_034013.1: (∼7 Mbp), Chr 6=NC_034014.1: (∼15 Mbp), Chr 7=NC_034015.1: (∼7 Mbp) and Chr 8=NC_034015.1: (∼10 Mbp). (**B**): chromosome wide LD decay of r^2^ (y-axis) over the physical distance in Mbp (x-axis). Each colored line represents a smoothed r2 for all marker pairs on each chromosome. The horizontal dashed red line indicates a cut-off r^2^=0.25.

To determine the extent of LD decay in the diversity panel, we estimated the pairwise LD coefficient (r^2^) at chromosomic level. LD decay was estimated for each chromosome by estimating the intersection of r^2^=0.25 with the physical distance (**Figure 2.B** and **Figure S2**). We found that LD dropped at short distance, ranging from 250 to 500 kbp along all chromosomes, with the exception of chromosome 5 (ca. 4.7 Mbp). After LD pruning, a total of 1,959 unlinked SNPs was kept for population structure and kinship estimations.

### Population structure

PCA analysis separated the germplasm panel into 4 sub-populations based on the genetic origin (landrace vs modern breeding line) and fruit type (peach vs nectarine) (**Figure S3)**. Clade 1 on the top left corner, grouped exclusively modern breeding lines of peach and nectarine. This group seems to be driven by the geographical origin as most of the accessions were originated from North America (**Table S1**). Clade 2 represents a diverse genetic entity gathering both landrace and breed peach varieties. Genotypes within this clade were originated from Spain and North America suggesting the presence of higher admixture that could arise due to the exchange of the germplasm material. In contrast, clades 3 and 4 contained only landrace peach accessions mostly from different regions of Spain, Europe and South Africa. A neighbor joining (NJ) tree also identified four clear clusters, as illustrated in **Figure S4.** Comparable results were obtained from fastSTRUCTURE and are provided in the GitHub repository.

### Critical evaluation of GWAS models

Genome wide association studies may be susceptible to bias in the presence of measurement errors. False positive and negative associations arising from population structure or/and family relatedness may lead to erroneous conclusions. The examination of Q-Q plots can be used as a straight visual inspection to determine the appropriate statistical method controlling the confounding effects. In fact, Q-Q plots illustrate the distribution of markers under the null hypothesis, by plotting the observed −log_10_ *P-*values (y-axis) versus the expected −log_10_ *P-*values (x-axis). If a sharp diagonal line is observed then the null hypothesis is respected and no significant associations are reported. However, an upper deviated tail from the diagonal line would likely indicate true associations. Upward inflation close to the line’s origin indicates suspicious false positives while downward deflated tail suggests false negatives.

We empirically evaluated the adjustment of seven models to our data and in **Figure 3**, we plot their Q-Q behavior for significantly associated traits. Despite yielding statistically significant associations, represented as bigger dots, both single locus models GLM and SUPER exhibited prominent inflation beyond the expected null line. This deviation starting close to the origin indicates false positive predictions due to confounding effects (population stratification or genotype relatedness). MLM and CMLM multi-locus models showed matching *P*-value distributions, therefore their Q-Q plots were overlaid. Except for harvest date, where the null hypothesis cannot be rejected with neither inflated nor deflated *P*-values, MLM and CMLM unveiled downshifted line tails when assessed with the rest of traits. Such a result may indicate that these tests were able to reduce false positive associations, but likely yielded false negative ones. Another complex model (MLMM) was found to follow the null hypothesis with both harvest date and flavonoids; nonetheless a slightly downward tail was discerned for fruit weight and sorbitol content. Although being the best-fitting model yielding marker-trait associations with harvest date and flavonoids, FarmCPU did not show the same statistical power with other traits. Finally, the observed *P*-values produced by Blink (green color) were lying on the diagonal line with clear deviated tails toward the y-axis for all six aforementioned traits. All in all, Blink seems to be the best calibrated model, appropriately controlling false positive and false negative effects. For these reasons, we consider Blink as the most suitable model, best adjusted with all phenotypic data and from here on the GWAS results are based on it.

**Figure 3.**
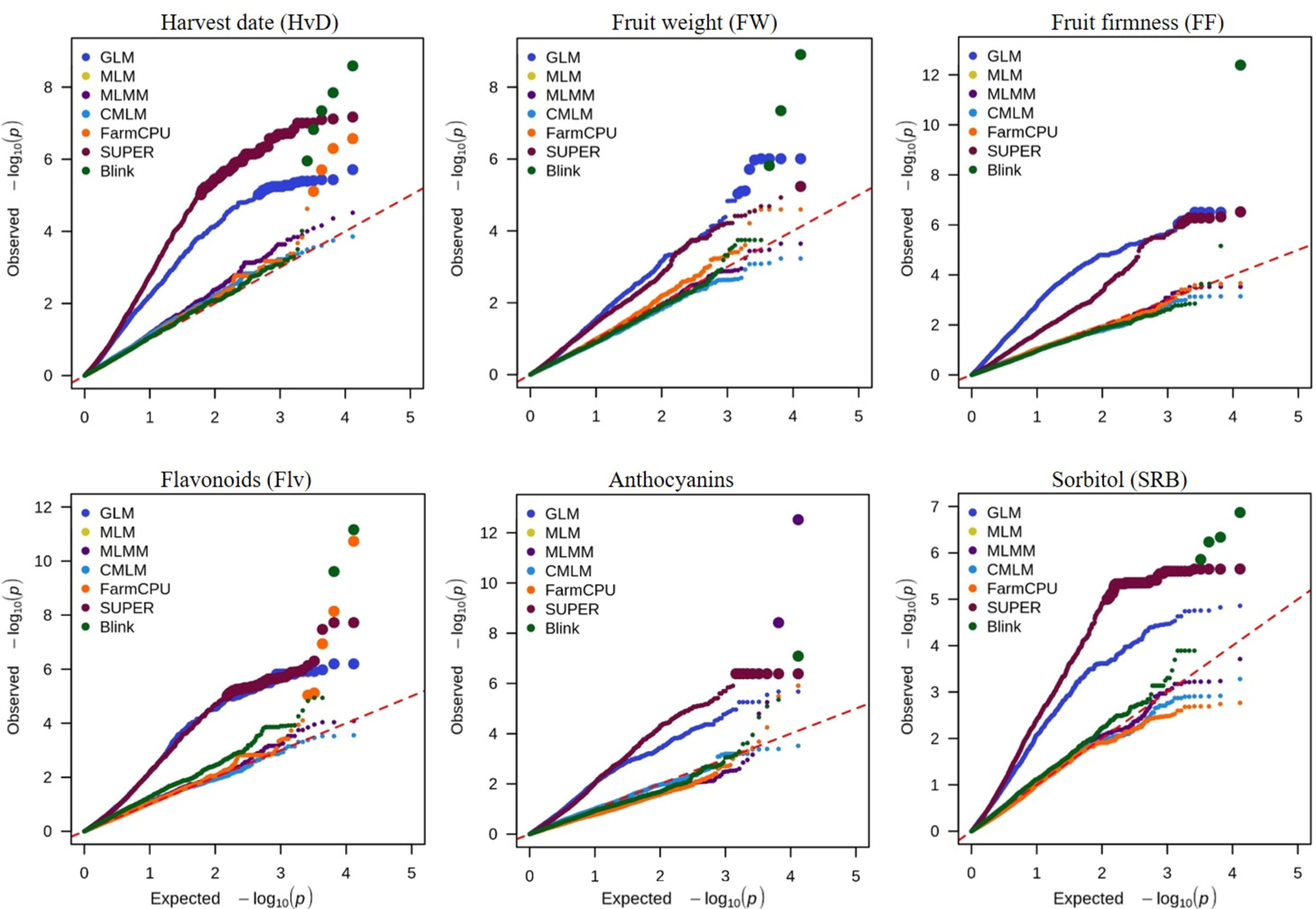
Q-Qplot comparison between the GWAS models implemented in GAPIT: General Linear Model (GLM), Mixed Linear Model (MLM), Compressed MLM (CMLM), Settlement of MLM under Progressively Exclusive Relationship (SUPER), Multiple Loci Mixed Linear Model (MLMM), Fixed and random model Circulating Probability Unification (FarmCPU) and Bayesian-information and Linkage-disequilibrium Iteratively nested keyway (BLINK). Note that MLM and CMLM models are overlaid. For each SNP, the expected −log_10_ transformed *P*-value (x-axis) is plotted against the −log_10_ the observed *P*-value (y-axis). The red dashed diagonal line corresponds to the expected Q-Q trendline under the null hypothesis (no association with the phenotype). Larger size dots refer to SNPs statistically associated with a trait. For clarity, only phenotypic traits with significant associations were represented.

### Marker-trait associations and identification of candidate genes

GWAS analysis was conducted on phenotypic traits with moderate to high heritability (H^2^ > 0.5). Consequently, contents of glucose, fructose, sucrose and total sugars were discarded from the subsequent analysis. To sum it up, among the remaining 12 traits, only six were found to be potentially influenced by polymorphic markers. Sixteen marker-trait association peaks were scattered throughout all chromosomes except chr 7 (**Table 2**). In the following sections we will discuss the results for each of these traits, namely harvest date, fruit weight, flesh firmness, and contents of flavonoids, anthocyanins, and sorbitol. For ease of interpretation, in the following paragraphs we summarize the lead SNPs and their corresponding LD blocks. The annotation of 250 kbp regions centering the peak SNPs resulted in a list of candidate causal genes provided in **Table S2**.

**Table 2.**
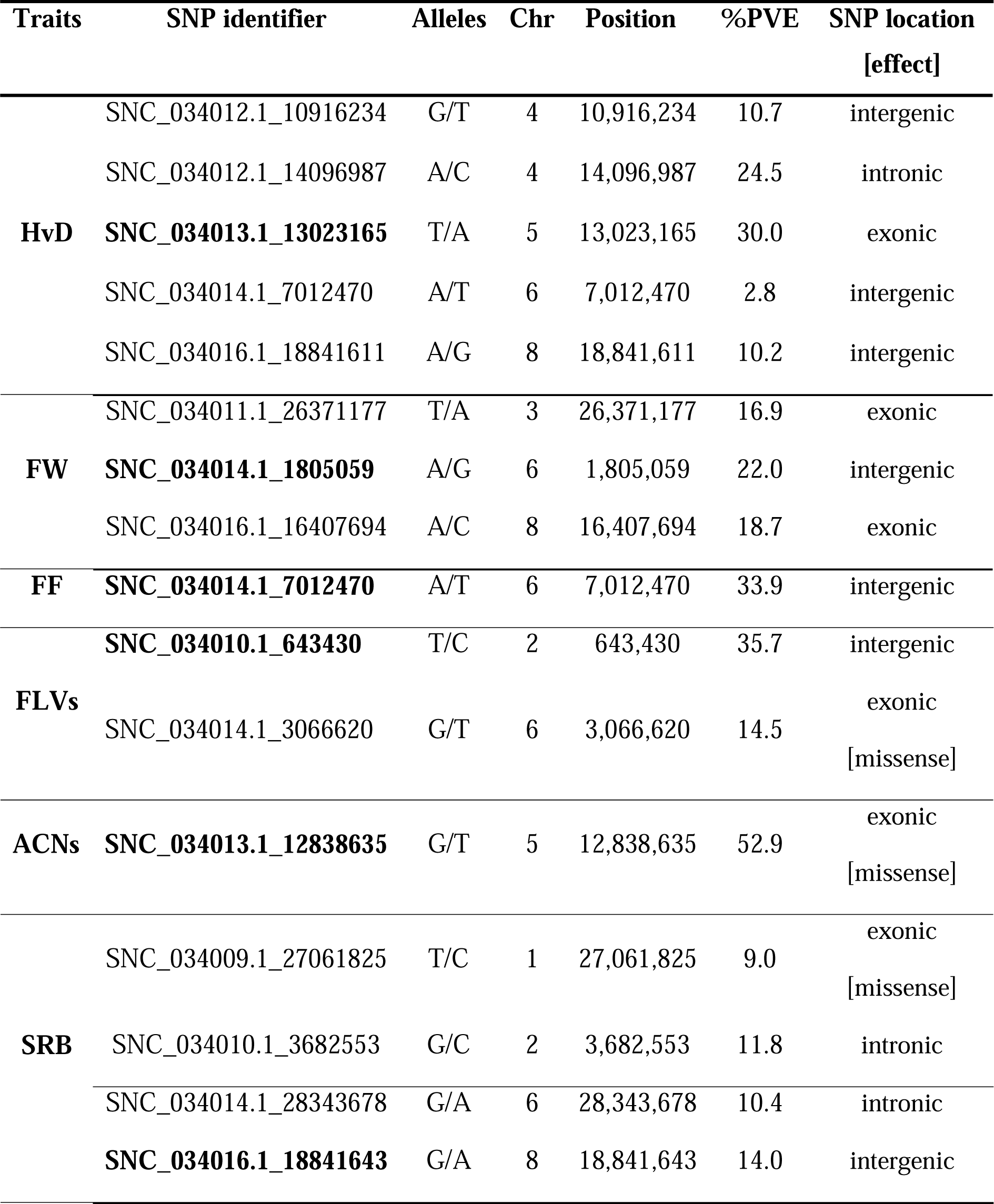
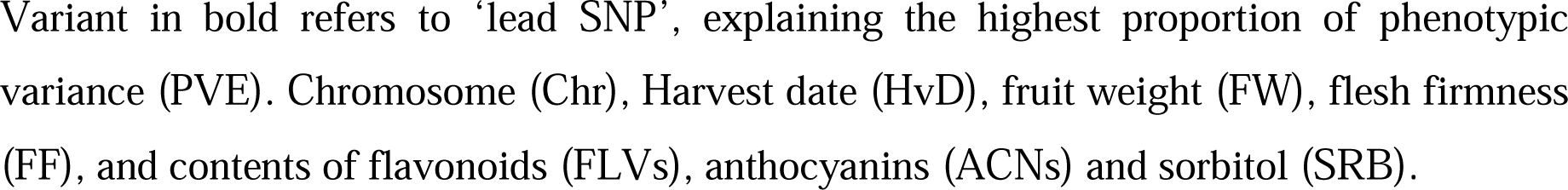
Information on significantly associated SNP markers with fruit-related traits in *Prunus persica.* Alleles are shown on the forward strand as reference/alternate.

### Harvest date (HvD)

The GWAS analysis resulted in five SNPs meeting the Bonferroni-adjusted threshold (**Figure 4**). Two SNPs were located on chr 4 and tagged as (SNC_034012.1_10916234, G/T) and (SNC_034012.1_14096987, A/C). Their allelic effect is summarized in **Figure S5**, where it can be seen that the first one correlates with delayed harvest and the second one with early one. Another associated marker was located on chr 5 (SNC_034013.1_13023165, T/A). Although covering the highest portion of %PVE, no significant allelic effect was observed (**Table 2**). This lead SNP was mapped within the first exon of *Prupe*.*5G138500*, a gene encoding a germin-like protein. One more significant site was identified on chr 6 and labeled as (SNC_034014.1_7012470, A/T). Allelic effect on phenotypic variation highlighted that both heterozygous and homozygous genotypes carrying the alternate allele (T) were lately harvested with respectively 6 and 13-days of delay (**Figure 4.C**). Similarly, the intergenic SNP located on chr 8 (SNC_034016.1_18841611, A/G), showed approximately 20-days delay in harvest date with heterozygous accessions (**Figure S5**).

**Figure 4.**
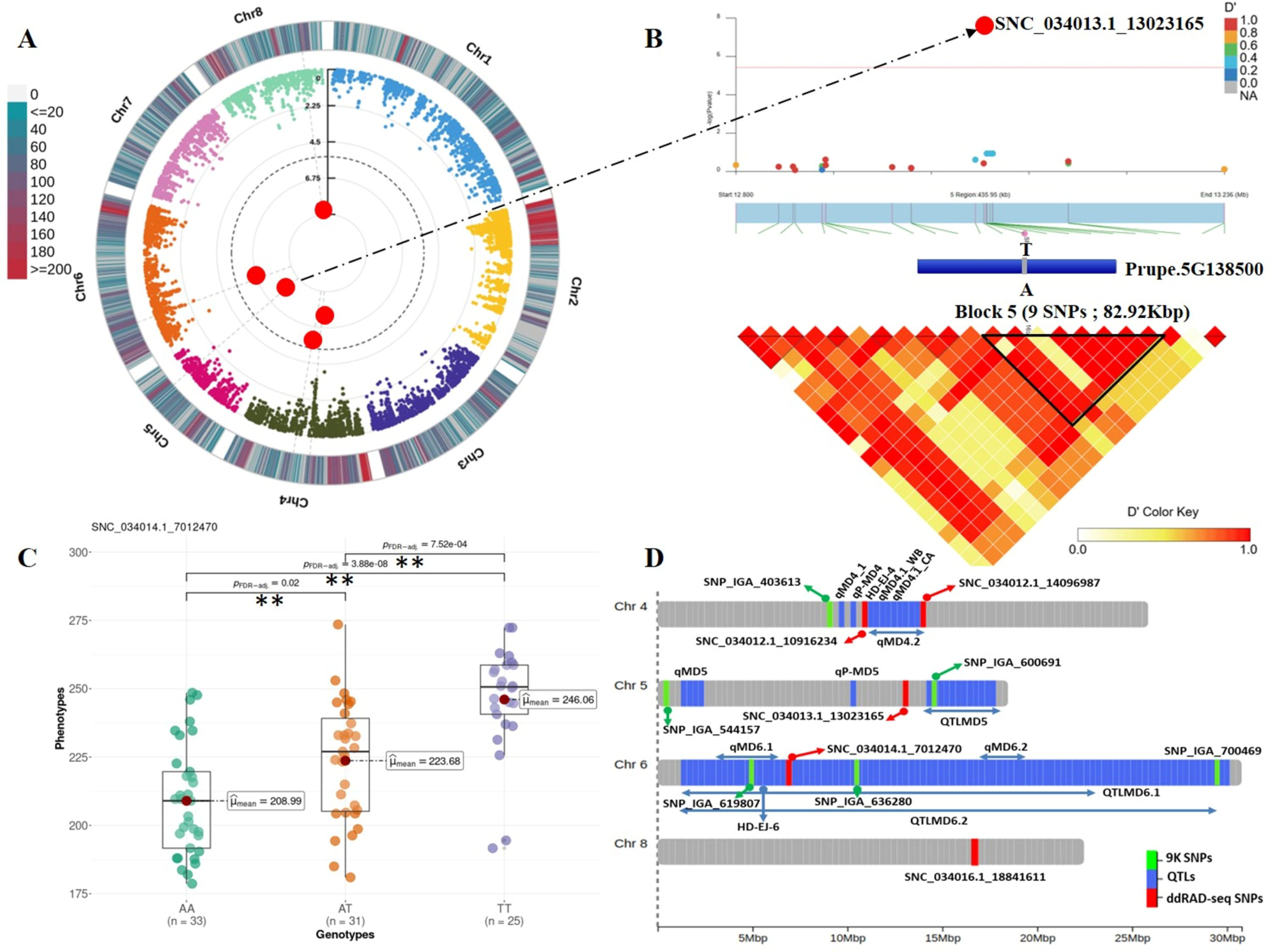
Genome Wide Association and LD block analysis for harvest date (HvD). (**A**): Circular Manhattan plot and association signals based on Blink model. Black dashed circular line corresponds to the Bonferroni adjusted threshold (-log_10_(*P*)=5.42). Red and large size dots correspond to statistically associated SNPs. Degradation from blue to red indicates the SNP density per 1 Mbp window across peach chromosomes. (**B**): Locus-specific Manhattan plot (upper panel) and LD heatmap (bottom panel) within 250 Kbp on either side of the lead SNP (SNC_034013.1_13023165). The prime candidate gene is represented as a blue box which in this case contains a single coding exon, where blue fragment refers to the exon. Pairwise LD measurements are displayed as D’ values with a color transition from yellow to red. (**C**): Boxplot depicting allelic effect of significant SNP on trait variation. Herein we highlight the SNP commonly affected Harvest date and fruit firmness. Mean value for each genotype is indicated by red circle and ** indicates significant pairwise comparison calculated by Games Howel test (*P* ≤ 0.05). (**D**): Genomic distribution of significant ddRAD-derived SNPs (red), reviewed QTLs in the literature (blue) and 9K array derived SNPs (green).

LD block analysis revealed various candidate genes, including cell wall modification (*Prupe*.*8G197700*: galacturonosyltransferase and *Prupe*.*8G199700*: cell division control protein), cytochrome P450 enzymes (*Prupe*.*8G196800*, *Prupe*.*8G196900*, *Prupe*.*8G197100* and *Prupe*.*8G197300*), UV-photoreceptor (*Prupe*.*4G185200*) and ethylene-responsive transcription factor (*Prupe*.*8G198700*).

### Fruit weight (FW)

Significant marker-trait associations were detected on three chromosomes: chr 3 (SNC_034011.1_26371177, T/A), chr 6 (SNC_034014.1_1805059, A/G) and chr 8 (SNC_034016.1_16407694, A/C). The explained variance oscillated between 17 and 22%, with SNC_034014.1_1805059 tagged as the lead intergenic marker (**Table 2**). The allelic effect of this lead marker (A/G) was found to be unfavorable, with the allele G associated with weight loss (∼22 grams) in homozygous accessions (**Figure 5.C**). A similar negative effect was observed with the SNP on chr 3 (T/A), with a significant reduction in fruit weight of 53g. Only marker mapped on chr 8 (A/C) was found to have a positive effect in heterozygous (**Figure S6**). Based on the LD block results, the lead SNP fell within the fourth block, a small interval (84 bp) overlapping no genes (**Figure 5.B**). Nonetheless, the associated SNPs did overlap protein-coding genes. Among them, genes encoding β-galactosidase (*Prupe*.*3G298200*), α-galactosyltransferase (*Prupe*.*3G298800*), thymidylate kinase (*Prupe*.*3G301400*) and transcription factors (GTE8: *Prupe*.*3G301300* and trihelix GT-4: *Prupe*.*3G300500*) (**Table S2**).

**Figure 5.**
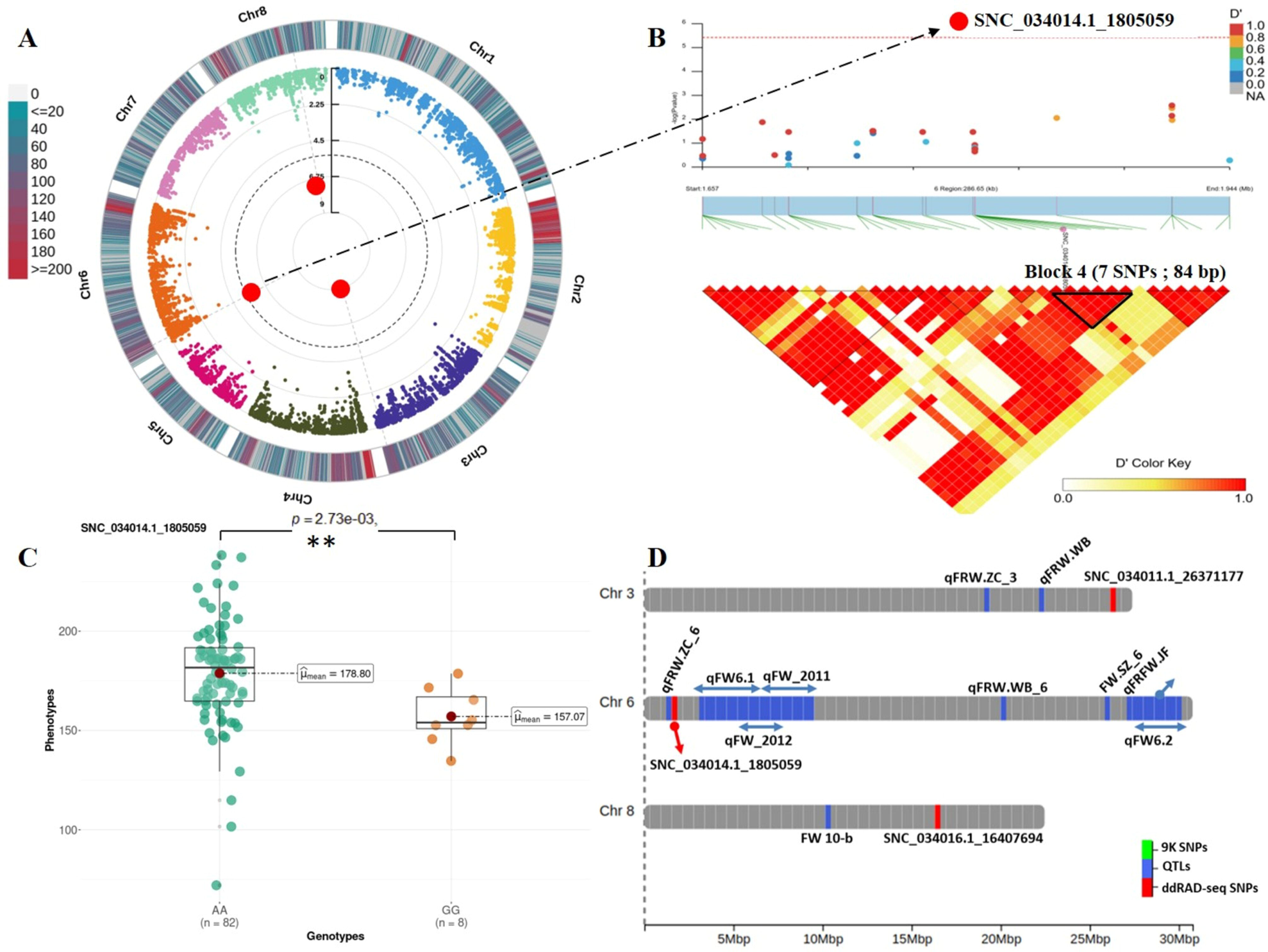
Genome Wide Association and LD block analysis for fruit weight (FW). (**A**): Circular Manhattan plot and association signals based on Blink model. Black dashed circular line corresponds to the Bonferroni adjusted threshold (-log_10_(*P*)=5.42). Red and large size dots correspond to statistically associated SNPs. Degradation from blue to red indicates the SNP density per 1 Mbp window across peach chromosomes. (**B**): Locus-specific Manhattan plot (upper panel) and LD heatmap (bottom panel) within 250 Kbp on either side of the lead SNP. Pairwise LD measurements are displayed as D’ values with a color transition from yellow to red. (**C**): Boxplot depicting allelic effect of lead SNP on trait variation. Mean value for each genotype is indicated by red circle and ** indicates significant pairwise comparisons calculated by Games Howel test (*P* ≤ 0.05). (**D**): Genomic distribution of significant ddRAD-derived SNPs (red) and reviewed QTLs in the literature (blue).

### Flesh Firmness (FF)

A single intergenic marker (SNC_034014.1_7012470; A/T) detected on chr 6 was statistically linked to flesh firmness and explained 33.9% of the total phenotypic variance (**Table 2**). This polymorphism showed a significant increase in the fruit firmness in both heterozygous and alternate homozygous genotypes which underlined the favorable effect of the alternative allele (T) on fruit firmness (**Figure 6.C**). It’s noteworthy to mention that this is the only marker simultaneously associated with two different traits (HvD and FF). Moreover, peach accessions carrying the aforementioned allele (either homozygous or heterozygous), were denoted late-harvested and firm peach accessions. Such a result may justify the high correlation existing between both traits (**Figure 1.B**).

**Figure 6.**
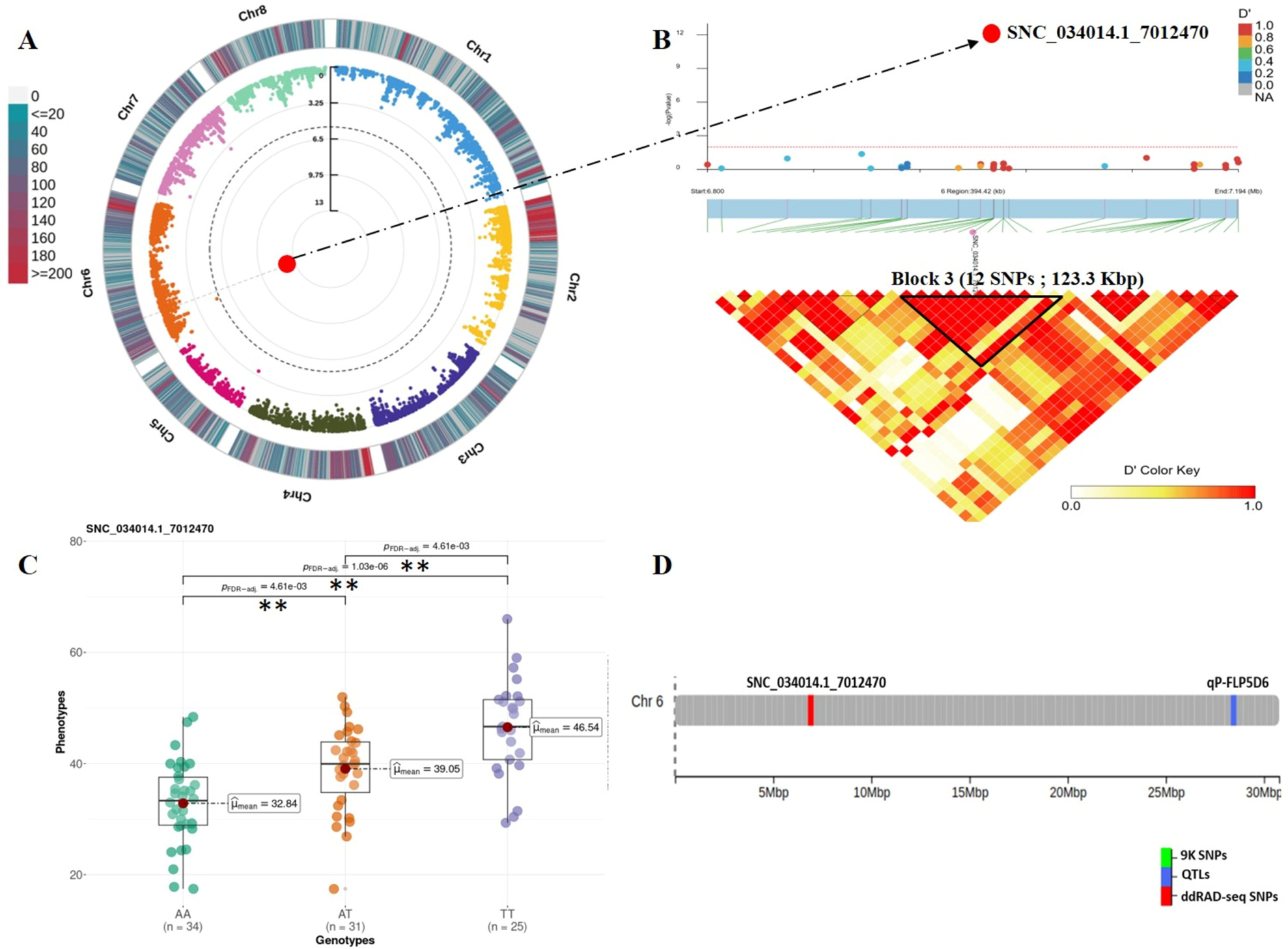
Genome Wide Association and LD block analysis for flesh firmness (FF). (**A**): Circular Manhattan plot and association signals based on Blink model. Black dashed circular line corresponds to the Bonferroni adjusted threshold (-log_10_(*P*)=5.42). Red and large size dots correspond to statistically associated SNPs. Degradation from blue to red indicates the SNP density per 1 Mbp window across peach chromosomes. (**B**): Locus-specific Manhattan plot (upper panel) and LD heatmap (bottom panel) within 250 Kbp on either side of the lead SNP. Pairwise LD measurements are displayed as D’ values with a color transition from yellow to red. (**C**): Boxplot depicting allelic effect of lead SNP on trait variation. Mean value for each genotype is indicated by red circle and ** indicates significant pairwise comparisons calculated by Games Howel test (*P* ≤ 0.05). (**D**): Genomic distribution of significant ddRAD-derived SNPs (red) and reviewed QTLs in the literature (blue)

By examining 250 kbp upstream and downstream the lead marker, it was found to reside in block 3, which makes it a relevant region to seek for candidate firmness-related genes. On the basis of their functional annotation, six genes were selected as potential candidates, including *Prupe.6G100500* encoding an E3 ubiquitin-protein ligase, *Prupe.6G101100* corresponding to vegetative cell wall protein, *Prupe*.*6G101600* annotated as aquaporin PIP2 and *Prupe.6G102300* encoding homeobox-leucine zipper transcription factor **(Table S2).**

### Flavonoids (Flvs)

The Manhattan plot displayed two peaks statistically associated with flavonoids content (**Figure S7.A**). The first peak was identified within the intergenic region of chr 2 and named as (SNC_034010.1_643430, T/C). The alternative allele (C) was marked as favorable for heterozygous (TC) and homozygous alternate (CC) genotypes since they showed approximately two-fold increase in the flavonoids content (**Figure S7.C**). The second associated SNP (SNC_034014.1_3066620; G/T) was located on chr 6 and physically mapped on the first exon of *Prupe.6G041500*; a candidate gene encoding a non-specific lipid-transfer protein-like (**Table S2**). The average flavonoids content in alternative homozygous peach accessions (TT) was significantly enhanced compared to the reference homozygous individuals (GG) (**Figure S8**). Thus, the T allele can be considered as a favorable one. Based on LD block results, we annotated a total of 14 genes (**Table S2**). According to their biological function and tissue-specific expression, we narrowed the list to a few promising ones, including two genes encoding transcription factors (*Prupe*.*2G009100*, bHLH and *Prupe*.*6G041400*, bZIP).

### Anthocyanins (ACNs)

Regarding the anthocyanins content, we detected a single peak signal on chr 5 exceeding the threshold line (**Figure S9.A**). This locus tagged as (SNC_034013.1_12838635; G/T) falls within exon 2 of *Prupe*.*5G134900*, encoding a B3 domain-containing transcription factor. Thus, *Prupe*.*5G134900* was considered as a prime candidate gene. The identified marker explained a large portion of the variation (53%), and was found to exert an unfavorable effect on anthocyanins content (**Figure S9.C**). Indeed, pairwise comparisons of SNP allelic effect showed a significantly lower anthocyanins content in the homozygous alternate individuals (TT) compared to the reference homozygous (GG). Screening for genes residing within LD block resulted in three further candidate genes involved in different biological functions (*Prupe.5G134200*, *Prupe*.*5G134800* and *Prupe.5G135200*) (**Table S2**).

### Sorbitol (SRB)

Four significant association signals dispersed on different chromosomes were predicted to affect the sorbitol content (**Table 2** and **Figure S10**). On chr 1, an intergenic SNP (SNC_034009.1_2706825; T/C) explained the lowest proportion of phenotypic variation. The SNP on chr 2 (SNC_034010.1_3682553; G/C), in the third intron of a gene encoding a flowering time control protein (*Prupe*.*2G0303400*), explained 12% of the PVE. Similarly, (SNC_034014.1_28343678; G/A) was located on chr 6 and mapped on the intronic region of *Prupe.6G320000*, a gene encoding a serine/arginine rich factor. Both *Prupe*.*2G0303400* and *Prupe*.*6G320000* are suggested as plausible sorbitol-related genes. The lead SNP explaining the highest PVE (14%) was identified in an intergenic region of chr 8 (SNC_034016.1_18841643; G/A).

With the exception of (SNC_034014.1_28343678) the remaining loci were observed to have desirable effect on sorbitol content (**Figure S11**). We identified 26 genes distributed in 250 kbp on either side of each associated SNP. Among them, some were discovered to be over-expressed in the fruit (Log_2_FC > L3L), including genes encoding heavy metal-associated isoprenylated proteins (*Prupe*.*2G033600*, *Prupe*.*2G033700* and *Prupe.6G321400*), pectinesterases (*Prupe*.*6G318500*), exonucleases (*Prupe*.*6G316100*), dormancy-associated proteins (*Prupe.6G319600*), cell cycle checkpoint control proteins (*Prupe*.*6G321300*) and the E3 ubiquitin-protein ligase RNF4 (*Prupe*.*8G199600*). A cluster of four cytochrome P450 encoding genes was also identified. This plethora of genes may shed light on several key processes that are subject to influence the sorbitol biosynthesis.

## Discussion

### Performance of variant callers

SNPs discovery in plant genomes has been a widely used strategy for developing molecular markers useful for MAS, genomic selection, phylogenetic analysis, etc. In order to detect and track these genetic variations, we performed a SNP discovery pipeline on paired-end reads mapped to a diploid genome using BCFtools, Freebayes, and GATK-HaplotypeCaller. SNP calling is known to be error prone. Spurious variants may have several sources; errors associated with sample processing (library preparation, PCR amplification), sequencing, as well as, computational analysis^19^. To remove likely false positive variants, best practices and carefully chosen cut-offs are needed. In our analysis, a SNP site was kept when passing the following filters: mapping and call quality, read depth, as well as call rate and MAF. Though either calling tool can be adapted, we observed a certain inconsistency in the number of high-quality SNPs revealed by each tool. Notably, GATK-HC exhibited the highest sensitivity in SNPs calling, followed by Freebayes then BCFtools. The outperformance of GATK-HC is actually not surprising as it heavily relies on local *de-novo* assembly of haplotypes in active regions^20^. In other terms and unlike the rest of tools, whenever GATK encounters regions with substantial evidence of variation relative to the reference, it discards the existing mapping information and reassembles the read mappings. Our results are in line with^21^ concluding that in *Arabidopsis thaliana,* GATK-HC was found to be more accurate compared to BCFtools. Additionally, GATK-HC had the lowest proportion of false positives compared to both Freebayes and BCFtools^22^. On the other hand, the variation in the number of detected SNPs may be partly due to the underlying algorithms. Indeed, GATK-HC and Freebayes are Bayesian variant detectors while BCFtools mpileup uses Hidden Markov Models. Although having an extensive format requirement (e.g: read group specified in the input header), GATK-HC seems to be more precise dealing with ddRAD-seq mapped reads in peach. Nevertheless, to further increase confidence, in this study we only considered SNPs called by all three approaches.

### Statistical model selection

Choosing a statistically reliable model is another fundamental pillar for a successful GWAS. Population structure and genetic relatedness are confounding factors increasing the rate of ambiguous associations and decreasing the statistical power. When ignored, they lead to substantial inflation of *P*-values as highlighted in the GLM model (**Figure 3**). In spite of including PCA components and kinship as covariates, SUPER model had also a large number of false positives. This may be explained by the fact that both GLM and SUPER are single-locus approaches failing to catch true associations when dissecting complex traits. Comparable inflated *P*-values were observed in *Arabidopsis thaliana* when testing flowering time, a polygenic trait, with the naïve model (GLM)^23^. In contrast, two other single-locus models, MLM and its compressed version (CMLM), were observed to adjust for false positives at the cost of failing to find any significant marker. Similar results were observed with MLMM, a multi-locus extension of MLM model (**Figure 3**). Overall, we conclude that MLM-based methods are likely missing potentially important SNPs.

The inspection of Q-Q plots declared FarmCPU and Blink as the most sophisticated algorithms yielding significant associations. Whereas FarmCPU returned significant signatures with only two traits (HvD and Flvs), Blink consistently inferred associations with six traits (HvD, FW, FF, Flvs, ACNs and SRB). FarmCPU and Blink have emerged to prevent over-fitting and to control false positives simultaneously^24, 25^. FarmCPU employs iteratively the fixed-effect model (FEM) and random effect model (REM) to eliminate confounding factors. FEM contains testing markers, one at a time, and associated markers as covariates to control false positives. To circumvent model over-fitting in FEM, the associated markers are estimated in REM and are used to derive the kinship^24^. Additionally, FarmCPU relies on the binning approach, where the whole genome is equally divided into bins and only the most significant marker is selected from each bin^24^. Despite its promise, this model is hampered by two major pitfalls: REM is computationally demanding and the assumption of bins rarely occurs in practice. As a consequence, Blink was designed to optimize the computational burden by substituting the REM with FEM through approximating maximum likelihood using the Bayesian Information Criterion and by increasing the statistical power by replacing the bin approach with the LD method^25^.

Overall, Blink seems to be the well-suited model for our set of data, balancing false positives and false negatives. This statement is underpinned by the GAPIT team, which already stated that Blink is statistically more powerful than FarmCPU^26^.

### Marker-trait association for the target traits

Out of 16 studied traits, association mapping using ddRAD-derived-SNPs and Blink, revealed association signals with six traits. Totally, 16 significant loci were inferred and distributed as follows: harvest date (chr 4, 5, 6 and 8), fruit weight (chr 3, 6 and 8), flesh firmness (chr 6), flavonoids (chr 2 and 6), anthocyanins (chr 5), and sorbitol (chr 1, 2, 6, and 8). Promising candidate genes were selected when residing within the LD block containing the significant loci, known to be related to the targeted trait and being over-expressed in fruit tissue. Our results were further discussed in comparison with^16^ which studied the same phenotypic data and germplasm material, but genotyped using the 9K SNP array instead.

### Harvest date

Peaches and nectarines are generally harvested at physiological maturity, then ripening off the trees. Harvest date and maturity date are frequently used as synonyms and are expressed in Julian days. HvD is defined as the day on which a certain percentage of peaches reach maturity. Maturity date (MD) is defined as the interval of time from the first day of the calendar year till the harvest date^27^. In our study, five association signals for HvD were highlighted. Two were inferred on chromosome 4 and the rest were distributed on chromosomes 5, 6 and 8.

As established by^28, 29^, major QTLs controlling maturity date have been reported on linkage groups LG4 and LG6 (**Table S3**). Particularly, a major QTL on LG4 referred to as qMD4.1 showed a pleiotropic effect on fruit weight and firmness^28, 30^. Interestingly, our marker SNC_034012.1_10916234 mapped at (∼10.91 Mbp), was overlapping the (qMD4.1_CA) locus from C×A progeny spanning the interval between 10.87-12.09 Mbp^30^. This same QTL from W×By progeny (qMD4.1_WB) was found 65 kbp from our marker (**Figure 4.D**). In the same vein, SNC_034012.1_10916234 was delimited by one downstream (HD-EJ-4)^31^ and two upstream quantitative loci (qP-MD4)^32^ and (qMD4_1)^33^ mapped respectively at 0.5, 5.3 and 1245 kbp from the SNP’s coordinate (**Figure 4.D**). Likewise, the second marker on chr 4 (SNC_034012.1_14096987) mapped at (∼14.09 Mbp) was found within the genomic region of strong confidence QTL (qMD4_2) spanning the interval (11.20 - 14.10 Mbp)^34^. Contrasting with associated SNP from the 9K assay^16^, our markers seems to be more confident as they are located within the QTL boundaries which supports their reliability. Altogether, we anticipate that the aforementioned SNPs on chr 4 could be integrated as promising markers for HvD breeding goals. As well, we conclude that LG4 seems to be a chromosomal hotspot hosting a cluster of major QTLs associated with the maturity date. QTLs influencing maturity date were also detected on LG4 in peach-related species, for instance; sweet cherry^35^. Therefore, we believe that this trait could be controlled by orthologous loci within *Prunus* species.

Marker ‘SNC_034013.1_13023165’ mapped on chr 5 (∼13.02 Mbp) was supported by an adjacent locus (QTLMD5) spanning the region (14.38 - 17.64 Mbp)^36^ and other distant signals (qP-MD5 and qMD5)^32, 34^. Significant markers from 9K array^16^ were found to be physically closer to the QTLs (**Figure 4.D** and **Table S3**). Finally, the significant SNP on chr 6 ‘SNC_034014.1_7012470’ was residing within two QTL intervals^36^ QTLMD6.1 and QTLMD6.1, supporting it. Similar findings were observed with 9K-associated markers.

Multiple candidate genes potentially influencing the harvest date were shortlisted (**Table S2**). Most importantly, an ethylene-responsive transcription factor (*Prupe*.*8G198700*). Ethylene-responsive elements are relevant in climacteric fruits and have been proposed as candidate genes for fruit maturation date in different *Prunus* species^31, 37^. We also identified a cell wall remodeling gene encoding galacturonosyltransferase. This finding is in consonance with^37^ defining a galacturonosyltransferase as a candidate gene for late harvested cultivars.

### Fruit weight

Fruit weight is a quantitative trait with great importance in peach breeding. Previous studies in peach have divulged that FW is monitored by multiple QTLs distributed across all chromosomes^34, 38–40^. Using GWAS, we identified a significant SNP on chr 3 (∼26.37 Mbp) located respectively at 4.07 and 7.27 Mbp downstream of two QTLs qFRW.ZC_3 and qFRW.WB (**Figure 5, Table S3**). On chr 6, another significant marker was predicted at (∼1.80 Mbp). This marker was delimited in near proximity by two reliable QTLs (qFRW.ZC_6)^40^ and (qFW6.1)^34^, situated respectively at 387 and 1,358 bp. On chr 8, SNC_034016.1_16407694, was localized at (∼6 Mbp) downstream of marker flanking QTL (FW 10-b)^39^. This is in contrast with^16^ where no associated loci were reported for this trait (**Table S3**). Such results support the relevance of our findings in dissecting the genetic control of complex fruit traits and shed light on the effectiveness of ddRAD-seq genotyping on inferring *novel* association signatures.

Candidate genes prediction revealed two transcription factors, trihelix GT-4 (*Prupe.3G300500*) and GTE-8 (*Prupe.3G301300*). Transcriptional regulators are abundant in plant genomes and they are implicated in various biological processes. Interestingly, trihelix genes are known to be photo-responsive proteins^41^. It’s well documented that light exposure affects fruit size, shape and quality^42^. Thus, we speculate that trihelix TF may regulate the fruit weight in peaches. Moreover, cell wall enzymes such as β-galactosidase, α-galactosyltransferase may act as key components of cell wall turnover during stone fruit growth^43^. Finally, thymidylate kinase exhibited strong upregulation suggesting a possible role in peach fruit development as validated in rice, barley and maize^44^.

### Flesh firmness

Firmness is a key textural indicator of peach quality and directly influences their shelf life. In our study, we identified a single firmness related locus SNC_034014.1_7012470 on chr 6. In the same LG6, a firmness loss QTL (qP-FL5d6) was described (**Figure 6.D** and **Table S3**). Another stable QTL (qP-FF6.1^m^) was also detected over two years in related species, particularly in sweet cherry^35^. Using 9K inferred SNPs and MLM model^16^, no significant association signals were found.

Four genes were selected as strong candidates encoding: ubiquitin-protein ligase (*Prupe*.*6G100500*), vegetative cell protein (*Prupe*.*6G101100*), aquaporin PIP2 (*Prupe*.*6G101600*) homeobox-leucine zipper protein (*Prupe*.*6G102300*). E3 ligase genes were found to be differentially expressed in either melting flesh or stony hard fruit during the ripening^48^. Aquaporins are transmembrane water transporters and water uptake within fruit is highly related with fruit firmness^45^. Thus, aquaporins could play a key role in maintaining cell turgor in peach. Finally, homeobox-leucine zipper proteins were denoted as potential biomarkers for the ripening process in peach^46^.

### Flavonoid and anthocyanin contents

Flavonoids are major polyphenol compounds playing a central role in fruit color and flavor. Our analysis yielded two potential association signatures in chr 2 and 6. These results go along with^47^ affirming that the majority of lead SNPs linked with many flavonoid metabolites in peach were located on chr 2. Herein, SNC_034010.1_643430 was supported by two QTLs^39^ identified in Venus × Bigtop progeny and named as ‘FLV 10-a’ and ‘FLV 10-b’ (**Figure S7.D** and **Table S3**). It’s well documented that flavonoid biosynthesis is a complex pathway, transcriptionally regulated by members of Myb and bHLH families^48^. Although no Myb encoding gene was found in our analysis, a highly up-regulated bHLH-TF was inferred and may be considered as a promising candidate gene involved in flavonoid regulation.

Anthocyanins constitute an important group of plant pigments belonging to the flavonoid family. Their differential accumulation in peach results in the distinctive fruit and flesh color^48^. Although there is strong evidence that their biosynthesis is mainly regulated by a Myb10 transcription factor on LG3, many anthocyanin-related QTLs were identified on LG4, LG5, and LG6^34, 39, 40^. Our analysis detected a single lead marker on chr 5 accounting for ∼53% of the PVE. Thus, ‘SNC_034013.1_12838635’ may be a preferential target for an effective marker assisted selection. It was delineated on both downstream and upstream sides by (qANT)^39^, (qATCYN.ZC)^40^ and (qPSC5)^34^. When genotyped with the 9K array^16^, no associated markers were detected on LG5. Remarkably, our polymorphic marker was physically falling in the exonic region of *Prupe*.*5G134900*, a gene encoding a B3 domain-containing transcription factor. Although the functional relevance of this prime gene requires further validation, we hypothesize that the genetic control of anthocyanins may be driven by B3 DNA-binding protein. Curiously, for both anthocyanins and flavonoids, a B3 family transcription was selected as candidate gene (respectively *Prupe*.*5G134900* and *Prupe*.*6G041000*). This may be explained by the fact that anthocyanins are a class of water-soluble flavonoids. Thus, it’s plausible to hypothesize that genes involved in flavonoids and anthocyanins regulation are in coordination.

### Sorbitol

Sugar content is one of the most important quality traits perceived by the consumers. The sweetness intensity depends on the overall sugar amount brought by sucrose, glucose, fructose and sorbitol. These first three sugar types were discarded from our analysis as they didn’t meet the heritability cutoff. Regarding the sorbitol, association signatures were found in chr 1 (∼27.06 Mbp), chr 2 (∼3.68 Mbp), chr 6 (∼28.34 Mbp) and chr 8 (∼18.84 Mbp). Genetic mapping has been extensively carried out to identify key QTLs responsible for sorbitol biosynthesis. A reliable QTL (qSOR_1) was mapped on the upper region of LG1, nearly 17.5 Mb upstream of our associated marker (**Figure S10.D**). Compared to the 9K association study^18^, no significant association signal was detected on LG1 (**Table S3**). On chr 2, we were able to find an adjacent QTL supporting the accuracy of our results^49^. Indeed, qSOR_2 was positioned at ∼1.2 Mbp from our marker SNC_034010.1_3682553.

Finally, this work depicts ddRAD-seq genotyping as an efficient approach for SNPs detection and association studies. Akin to the 9K SNP array, ddRAD-seq yielded valuable markers strongly supported by stable QTLs. However, while SNP arrays are engineered to specifically include polymorphic loci from genomic regions of interest and focus on harboring SNPs known to be linked to commercially important traits, ddRAD-seq samples the genome randomly, without prior knowledge of target regions. For this reason, ddRAD-seq might be a better fit for analyses concerned with unexplored biological processes.

Concisely, we successfully used ddRAD-seq-derived SNPs to identify genomic regions and genes influencing major fruit-related traits in peaches. The inferred associated SNPs appeared to be reliable as they often explained a fairly high percentage of the total phenotypic variance. The survey of candidate genes for these relevant polymorphic sites rendered plenty of genes implicated in various processes. Genes harboring significant markers may be considered as preferential targets for peach breeding. However, due to the complexity of the examined traits, future functional validation would provide additional hints to support the breeding efforts.

## Material and Methods

### Plant material and phenotypic evaluation

A total of 90 peach and nectarine accessions were used for double digest restriction-site associated sequencing (ddRAD-seq) and subsequent GWAS analysis. The germplasm panel comprises 73 landraces and 17 modern breeding lines originating from Spain, United States, France, Italy, New Zealand, and South Africa. All genotypes were grown under Mediterranean soil conditions at the Experimental Station of Aula Dei (CSIC) located at Zaragoza, Spain (41.7245 °N, 0.8118 °W) and analyzed during three fruiting seasons (2008-2010). Information about plant accessions is summarized in **Table S1**.

The phenotypic data previously reported by^16^ were re-analyzed in the present study. Briefly, 16 traits were evaluated by randomly harvesting 20 fruits from each cultivar at the commercial maturity during three years. Traits were split into two categories. Agronomic features included harvest date (HvD; Julian days), fruit weight (FW; grams), flesh firmness (FF; Newton), soluble solids content (SSC; °Brix), titratable acidity (TA; grams malic acid/100 g flesh weight) and ripening index (RI; SSC/TA). Besides, biochemical variables comprised vitamin C (Vit C; mg of ascorbic acid/100 g flesh weight), total phenolics (Phen; mg of gallic acid equivalents/100 g flesh weight), contents of flavonoid (Flv; catechin equivalents/100 g flesh weight) and anthocyanin (ACNs; cyanidin-3-glucoside/kg flesh weight), sucrose (Suc; g/kg flesh weight), glucose (Glu; g/kg flesh weight), fructose (Fruc; g/kg flesh weight), sorbitol (SRB; g/kg flesh weight), and total sugars (TS; g/kg flesh weight) and relative antioxidant capacity (RAC; μg TE/g flesh weight).

Variance components and broad sense heritability (H^2^) were estimated using the variability R package v0.1.0. Only traits with H^2^ > 0.5 were considered for association analysis. Distribution of averaged phenotypic data was checked in R using Shapiro-Wilk test. Non-normal distributions were transformed using bestNormalize package (v1.8.3)^50^.

### DNA extraction and enzyme evaluation

Genomic DNA was extracted from leaves using the DNeasy Plant Mini Kit (Qiagen, Dusseldorf, Germany) following the manufacturer’s recommendations. DNA concentration and quality were checked using PicoGreen®dye and measured in a fluorospectrometer. Whole-genome genotyping was carried out using ddRAD-seq approach by combining low and high frequency cutter to digest DNA; respectively *Pst1* and *Mbol* as described in peach by^51^. This enzyme pair yielded the highest number of loci with a size range between 300 and 400 bp and prevented repetitive region sampling. Selected loci are those having the sticky ends of both enzymes^52^.

### ddRAD libraries preparation and sequencing

DNA libraries were constructed at the Genomic Unit at IABiMo INTA-CONICET (Argentina) following^51, 52^ recommendations. Shortly, digested DNA with *Pst1*/*Mbol* pair were gel excised, eluted then ligated to barcoded adapters specific to each sample. Ligated fragments from 24 samples were subsequently pooled together and were PCR amplified with indexed primers to tag each pool. Finally, paired-end reads (250 bp) were generated on an Illumina NovaSeq 6000 instrument at CIMMYT, Mexico. The raw sequencing data was deposited in the European Nucleotide Archive (ENA) under the BioProject PRJEB62784.

### Data processing and alignment

Raw reads were de-multiplexed and trimmed using the process-radtag module from STACKS suite (v2.59)^53^. After quality assessment, paired-end reads were mapped to *Prunus persica* reference genome v2 (GCF_000346465.2, retrieved from NCBI RefSeq^54^ using BWA-mem (v0.7.17)^55^. Redundant reads known as PCR duplicates were expurgated from downstream analysis as described by^22^. The resultant de-duplicated files were sorted and indexed using SAMtools^56^ to be ready for variant calling.

### Variant discovery pipeline

Variant calling was conducted in a single-sample mode testing the performance of three variant callers: BCFtools (v1.7)^56^, Freebayes (v1.0.0)^57^ and GATK-HaplotypeCaller (v4.2.3.0)^20^. Raw SNPs underwent standard quality filtering based on mapping quality (MQ > 40), variant quality (QUAL < 30) and depth of reads (DP ≥ 5) to remove artifactual calls. Consequently, clean SNPs from each calling method were merged by position and by reference/alternative alleles into multi-sample VCF files. SNPs resulting from the intersection of multi-samples VCFs were considered as highly accurate calls and were inspected to remove multi-allelic variants and those assigned to scaffolds. Then, they were filtered by call rate > 80% and residual missing genotypes were imputed with beagle’s default settings (v4.1)60. The imputation accuracy was evaluated in Tassel (v5.0)^58^ by masking 1% of the genotype and calculating the error rate. SNPs with (MAF > 0.05) were selected as a final call set to determine the population structure and marker-trait associations. SNP identifiers were created by concatenating their assigned chromosome and their base pair position (eg: SNC_034014.1_7012470).

### Linkage disequilibrium and population structure

Intra-chromosomal LD was calculated using Plink (v1.9)^59^, as a measure of *Pearson* correlation coefficient (r^2^) between marker-pairs. For each chromosome, LD distribution was plotted against its physical distance (Mbp). The LD decay curve was estimated as the average of r^2^ variation across 100 kbp bins and was fitted in R program. LD decay was defined by setting r^2^=0.25 as a threshold. LD decay extent was defined as the physical genomic distance at which the r^2^ decreased to half of its maximum value. Polymorphic sites showing strong LD were pruned in Plink by delimiting a window of 10 SNPs, removing one of the SNPs pair with r^2^ > 0.25 and then shifting the window 5 SNPs forward repeatedly. Genetic distance and kinship matrix between pairs of genotypes were computed using the centered identity-by-state method implemented in Tassel.

LD-pruned SNPs were selected to infer the population stratification of the GWAS panel using two complementary approaches. First, the Bayesian clustering algorithm implemented in fastSTRUCTURE (v1.04)^60^ was tested on predefined K subgroups ranging from 1 to 10. The optimal K value was estimated based on the lowest cross validation error. Then, principal component analysis was computed with SmartPCA (v1.1.0) R-package^61^.

### Genome wide association study

For association mapping, seven statistical models, ranging from single to multi-locus, were simultaneously tested in GAPIT (v3.1.0)^62^ Single locus models include general linear model (GLM), mixed linear model (MLM), compressed MLM (CMLM), and settlement of MLMs under progressively exclusive relationship (SUPER). Multi-locus algorithms comprise multiple loci mixed linear model (MLMM), fixed and random model circulating probability unification (FarmCPU), and Bayesian-information and linkage-disequilibrium iteratively nested keyway (Blink). Except for GLM, where no genotype relatedness was involved, population structure and kinship were both fitted as covariates in all models. Indeed, the first four PCA components and kinship were introduced respectively as fixed and random effects to reduce the false positives. The statistical model best fitting the data was chosen based on the quantile-quantile plot and the number of significant markers. Significantly associated markers were shortlisted based on the Bonferroni correction (-log_10_(0.05)/13045 = 5.42) and Manhattan plots were generated accordingly using CMplot package (v4.2.0)^63^. Statistically significant SNPs explaining at least 10% of the phenotypic variance (%PVE) were considered as most promising predictions and used for LD block analysis. Moreover, markers accounting for the largest proportion of phenotypic variance are hereinafter referred as ‘lead SNPs’.

### Annotation of SNP effects and identification of favorable alleles

First, genomic coordinates of SNPs were used to query Ensembl Plants REST services in order to obtain annotations of their effect on neighbor genomic features. In particular we used the Ensembl Variant Effect Predictor (VEP)^64^ and a modification of recipe R8^65^.

Then, allelic effect of significant SNP loci on trait variation was estimated through pairwise comparisons between the phenotypic values of the different genotypes: homozygous reference (0/0), heterozygous (0/1) and homozygous alternative (1/1). An allele is defined as favorable when a significant increase of the phenotypic value was observed between the homozygous reference and the remaining genotypes. Pairwise comparisons were run using the Games-Howell test and *P*-values were corrected for multiple testing using the FDR method. Results were visualized using ggstatsplot R-package (v0.9.1)^66^.

### LD-block analysis and identification of candidate genes

Significant SNPs were examined to identify candidate genes. Initially, it was considered whether polymorphisms would be localized in genic regions. Thereby, SNPs were mapped based on their physical position to *Prunus persica* genome (GCF_000346465.2). SNP-anchored genes were called ‘prime candidates’. Strong candidate genes were shortlisted when meeting three criteria: falling within the LD-block region harboring the significant SNPs, being functionally related to the trait of interest and being differentially expressed in fruit tissue. Expression information was retrieved from a recent study by^67^ which defined modules of co-expressed genes across different peach tissues and under various experimental conditions. Differentially expressed genes were those outlined in fruit experiments, particularly under cold storage and chilling injury.

LD-blocks were identified within 250 kbp windows upstream and downstream the lead sites. Block boundaries were delimited using a solid spine partitioning approach from LDBlockShow tool^68^. A block is defined as a group of SNPs that are in strong LD (D’ ≥ 0.7) with the first and last marker in the same block. A D’ value of 0 denotes complete linkage equilibrium, which implies frequent pairwise recombination between markers. Conversely, a D’ value of 1 indicates a complete linkage disequilibrium. Note that D’ and r^2^ are common measures of non-random association between two or more loci; while D’ refers to the co-inheritance of two alleles, r^2^ considers the allele frequency to distinguish between common and rare. Identified LD blocks were therefore scanned for candidate genes via NCBI genome data viewer^69^.

### QTLs review for fruit quality traits in peach

To benchmark the accuracy of our results, an exhaustive bibliographic review of previously reported QTLs mapped in the same linkage group as the associated markers was done. In a practical term, if an associated SNP is located nearby or within a QTL interval, then it’s considered as highly accurate and likely segregate with the observed trait. In case that the QTL boundaries are not defined as physical intervals (in bp), we used the nearest or the co-localizing markers as reference. Herein, we refer to the nearest marker as the closest one with a maximum of 5 cM from the QTL hotspot while the co-localizing marker is the one mapped within the QTL boundaries. Moreover, we calculated the physical distance separating our predicted associated markers from the QTLs. Finally, we compared these distances with a previous study using the same phenotypic data and peach material, although genotyped using the 9K SNP array^16^.

## Data availability

Raw sequence data and final variant call file (vcf) have been deposited in the European Nucleotide Archive (ENA) under the BioProject PRJEB62784 (data will be released at the publication date). Source code and documentation can be accessed at https://github.com/najlaksouri/GWAS-Workflow.

## Funding and Acknowledgments

This work was funded by Spanish Research Agency [grants AGL2014-52063-R, AGL2017-83358-R MCIN/AEI/10.13039/501100011033 and by “ERDF A way of making Europe”] and the Government of Aragón [grantsA09_20R, A09_23R, A10_20R], which were co-financed with FEDER funds and the CSIC [grant2020AEP119]. N.K. was funded with a pre-doctoral contract awarded by the Government of Aragón (2018-2023) and C.F-i-F. by a JAE-Pre fellowship from CSIC (2008-2013).

The authors wish to thank of personal of Genomic Unit at IABiMo INTA-CONICET (Argentina), in particular to A.F. Puebla for their technical assistance and for providing lab facilities; to R. Giménez, S. Segura, E. Sierra for their technical lab assistance and plant management in the field; and to M.A. Moreno for providing plant material.

## Contributions

Y.G., and B.C-M. conceived the project and its components. C.F-i-F. collected the samples, extracted DNA and performed the phenotyping. G.S helped to process genotyping. N.K performed the bioinformatic analysis. Y.G., and B.C-M. assisted the analysis and discussed the results. N.K wrote the manuscript. Y.G., and B.C-M. reviewed and edited the text. All authors read and approved the article.

## References

1. Micheletti, D. et al. Whole-genome analysis of diversity and SNP-major gene association in peach germplasm. PLoS One 10, 1–19 (2015).

2. Gogorcena, Y., Sánchez, G., Moreno-Vázquez, S., Pérez, S. & Ksouri, N. Genomic-based breeding for climate-smart peach varieties. in Genome designing of climate-smart fruit crops (ed. Kole, C.) 291–351 (Springer-Nature, 2020). doi:https://doi.org/10.1007/978-3-319-97946-5_9.

3. Cirilli, M. et al. Genetic and phenotypic analyses reveal major quantitative loci associated to fruit size and shape traits in a non-flat peach collection (*P. persica* L. Batsch). Hortic. Res. 8, 1–17 (2021).

4. Da Silva Linge, C., et al. Multi-locus genome-wide association studies reveal fruit quality hotspots in peach genome. Front. Plant Sci. 12, 1–18 (2021).

5. Aranzana, M. J., et al. *Prunus* genetics and applications after *de novo* genome sequencing: achievements and prospects. Hortic. Res. 6, 1–25 at https://doi.org/10.1038/s41438-019-0140-8 (2019).

6. De Mori, G. & Cipriani, G. Marker-assisted selection in breeding for fruit trait improvement: a review. Int. J. Mol. Sci. 24, 1–39 (2023).

7. Kumar, S. et al. GWAS provides new insights into the genetic mechanisms of phytochemicals production and red skin colour in apple. Hortic. Res. 9, 1–10 (2022).

8. Holušová, K. et al. High-resolution genome-wide association study of a large Czech collection of sweet cherry (*Prunus avium* L.) on fruit maturity and quality traits. Hortic. Res. 10, 1–15 (2023).

9. Pavan, S. et al. Recommendations for choosing the genotyping method and best practices for quality control in crop genome-wide association studies. Front. Genet. 11, (2020).

10. Verde, I. et al. Development and evaluation of a 9K SNP array for peach by internationally coordinated SNP detection and validation in breeding germplasm. PLoS One 7, e35668 (2012).

11. Thurow, L. B., Gasic, K., Bassols Raseira, M. do C., Bonow, S. & Marques Castro, C. Genome-wide SNP discovery through genotyping by sequencing, population structure, and linkage disequilibrium in Brazilian peach breeding germplasm. Tree Genet. Genomes 16, 1–14 (2020).

12. Mas-Gómez, J., Cantín, C. M., Moreno, M. Á. & Martínez-García, P. J. Genetic diversity and genome-wide association study of morphological and quality traits in peach using two Spanish peach germplasm collections. Front. Plant Sci. 13, 1–14 (2022).

13. Geibel, J. et al. How array design creates SNP ascertainment bias. PLoS One 16, 1– 23 (2021).

14. Peterson, B. K., Weber, J. N., Kay, E. H., Fisher, H. S. & Hoekstra, H. E. Double digest RADseq: An inexpensive method for *de novo* SNP discovery and genotyping in model and non-model species. PLoS One 7, 1–11 (2012).

15. Font i Forcada, C., Oraguzie, N., Igartua, E., Moreno, M. Á. & Gogorcena, Y. Population structure and marker-trait associations for pomological traits in peach and nectarine cultivars. Tree Genet. Genomes 9, 331–349 (2013).

16. Font i. Forcada, C., Guajardo, V., Chin-Wo, S. R. & Moreno, M. Á. Association mapping analysis for fruit quality traits in *Prunus persica* using SNP markers. Front. Plant Sci. 9, 1–12 (2019).

17. Cao, K. et al. Genome-wide association study of 12 agronomic traits in peach. Nat. Commun. 7, 1–10 (2016).

18. Verde, I. et al. The high-quality draft genome of peach (*Prunus persica*) identifies unique patterns of genetic diversity, domestication and genome evolution. Nat. Genet. 45, 487–494 (2013).

19. Olson, N. D. et al. Best practices for evaluating single nucleotide variant calling methods for microbial genomics. Front. Genet. 6, 1–15 (2015).

20. Van der Auwera, G. A. et al. From fastq data to high-confidence variant calls: The genome analysis toolkit best practices pipeline. Current Protocols in Bioinformatics (2013).

21. Schilbert, H. M., Rempel, A. & Pucker, B. Comparison of read mapping and variant calling tools for the analysis of plant NGS data. Plants 9, 1–14 (2020).

22. Ksouri, N. et al. ddRAD-seq variant calling in peach and the effect of removing PCR duplicates. Acta Hortic. 1352, 405–412 (2022).

23. Atwell, S. et al. Genome-wide association study of 107 phenotypes in *Arabidopsis thaliana* inbred lines. Nature 465, 627–631 (2010).

24. Liu, X., Huang, M., Fan, B., Buckler, E. S. & Zhang, Z. Iterative usage of fixed and random effect models for powerful and efficient genome-wide association studies. PLoS Genet. 12, 1–24 (2016).

25. Huang, M., Liu, X., Zhou, Y., Summers, R. M. & Zhang, Z. BLINK: A package for the next level of genome-wide association studies with both individuals and markers in the millions. Gigascience 8, 1–12 (2019).

26. Wang, J., Tang, Y. & Zhang, Z. Performing genome-wide association studies with multiple models using GAPIT. in Genome-Wide Association Studies (eds. Torkamaneh, D. & Belzile, F.) 199–217 (Springer US, 2022). doi:10.1007/978-1-0716-2237-7_13.

27. Veerappan, K., Natarajan, S., Chung, H. & Park, J. Molecular insights of fruit quality traits in peaches, Prunus persica. Plants 10, 1–14 (2021).

28. Eduardo, I. et al. QTL analysis of fruit quality traits in two peach intraspecific populations and importance of maturity date pleiotropic effect. Tree Genet. Genomes 7, 323–335 (2011).

29. Dirlewanger, E. et al. Comparison of the genetic determinism of two key phenological traits, flowering and maturity dates, in three *Prunus* species: Peach, apricot and sweet cherry. Heredity 109, 280–292 (2012).

30. Pirona, R. et al. Fine mapping and identification of a candidate gene for a major locus controlling maturity date in peach. BMC Plant Biol. 13, 1–13 (2013).

31. Romeu, J. F. et al. Quantitative trait loci affecting reproductive phenology in peach. BMC Plant Biol. 14, 1–16 (2014).

32. Serra, O. et al. Genetic analysis of the slow-melting flesh character in peach. Tree Genet. Genomes 13, 1–13 (2017).

33. Kalluri, N., Eduardo, I. & Arús, P. Comparative QTL analysis in peach ‘Earlygold’ F2 and backcross progenies. Sci. Hortic. 293, 1–8 (2022).

34. Hernández Mora, J. R., et al. Integrated QTL detection for key breeding traits in multiple peach progenies. BMC Genomics 18, 1–15 (2017).

35. Calle, A. & Wünsch, A. Multiple-population QTL mapping of maturity and fruit-quality traits reveals LG4 region as a breeding target in sweet cherry (*Prunus avium* L.). Hortic. Res. 7, 1–13 (2020).

36. Nuñez-Lillo, G. et al. High-density genetic map and QTL analysis of soluble solid content, maturity date, and mealiness in peach using genotyping by sequencing. Sci. Hortic. 257, 1–11 (2019).

37. Núñez-Lillo, G. et al. Transcriptome and gene regulatory network analyses reveal new transcription factors in mature fruit associated with harvest date in *Prunus persica*. Plants 11, 1–17 (2022).

38. Da Silva Linge, C., et al. Genetic dissection of fruit weight and size in an F2 peach (*Prunus persica* (L.) Batsch) progeny. Mol. Breed. 35, 1–19 (2015).

39. Zeballos, J. L. et al. Mapping QTLs associated with fruit quality traits in peach [*Prunus persica* (L.) Batsch] using SNP maps. Tree Genet. Genomes 12, 1–7 (2016).

40. Abdelghafar, A., Da Silva Linge, C., Okie, W. R. & Gasic, K. Mapping QTLs for phytochemical compounds and fruit quality in peach. Mol. Breed. 40, 1–18 (2020).

41. Kaplan-Levy, R. N., Brewer, P. B., Quon, T. & Smyth, D. R. The trihelix family of transcription factors - light, stress and development. Trends Plant Sci. 17, 163–171 (2012).

42. Reale, L. et al. The influence of light on olive (*Olea europaea* L.) fruit development is cultivar dependent. Front. Plant Sci. 10, 1–10 (2019).

43. Canton, M. et al. Metabolism of stone fruits: reciprocal contribution between primary metabolism and cell wall. Front. Plant Sci. 11, 1–10 (2020).

44. Zhu, L. et al. A shortest-path-based method for the analysis and prediction of fruit-related genes in Arabidopsis thaliana. PLoS One 11, 1–14 (2016).

45. Quirante-Moya, F., Martinez-Alonso, A., Lopez-Zaplana, A., Bárzana, G. & Carvajal, M. Water relations after Ca, B and Si application determine fruit physical quality in relation to aquaporins in *Prunus*. Sci. Hortic. 293, 1–14 (2022).

46. Nilo-Poyanco, R., Moraga, C., Benedetto, G., Orellana, A. & Almeida, A. M. Shotgun proteomics of peach fruit reveals major metabolic pathways associated to ripening. BMC Genomics 22, 1–29 (2021).

47. Cao, K. et al. Combined nature and human selections reshaped peach fruit metabolome. Genome Biol. 23, 1–25 (2022).

48. Wang, J. et al. Two MYB and three bHLH family genes participate in anthocyanin accumulation in the flesh of peach fruit treated with glucose, sucrose, sorbitol, and fructose *in vitro*. Plants 11, 1–14 (2022).

49. Desnoues, E. et al. Dynamic QTLs for sugars and enzyme activities provide an overview of genetic control of sugar metabolism during peach fruit development. J. Exp. Bot. 67, 3419–3431 (2016).

50. Peterson, R. A. Finding optimal normalizing transformations via bestNormalize. Contrib. Res. J. 13, 310–329 (2021).

51. Aballay, M. M., Aguirre, N. C., Filippi, C. V., Valentini, G. H. & Sánchez, G. Fine-tuning the performance of ddRAD-seq in the peach genome. Sci. Rep. 11, 1–13 (2021).

52. Aguirre, N. C. et al. Optimizing ddRADseq in non-model species: A case study in Eucalyptus dunnii Maiden. Agronomy 9, 1–21 (2019).

53. Rochette, N. C., Rivera-Colón, A. G. & Catchen, J. M. Stacks 2: Analytical methods for paired-end sequencing improve RADseq-based population genomics. Mol. Ecol. 28, 4737–4754 (2019).

54. O’Leary, N. A. et al. Reference sequence (RefSeq) database at NCBI: Current status, taxonomic expansion, and functional annotation. Nucleic Acids Res. 44, D733–D745 (2016).

55. Li, H. & Durbin, R. Fast and accurate short read alignment with Burrows-Wheeler transform. Bioinformatics 25, 1754–1760 (2009).

56. Danecek, P. et al. Twelve years of SAMtools and BCFtools. Gigascience 10, 1–4 (2021).

57. Garrison, E. & Marth, G. Haplotype-based variant detection from short-read sequencing. ArXiv 1207, 1–5 (2012).

58. Bradbury, P. J. et al. TASSEL: Software for association mapping of complex traits in diverse samples. Bioinformatics 23, 2633–2635 (2007).

59. Purcell, S. et al. PLINK: A tool set for whole-genome association and population-based linkage analyses. Am. J. Hum. Genet. 81, 559–575 (2007).

60. Raj, A., Stephens, M. & Pritchard, J. K. FastSTRUCTURE: Variational inference of population structure in large SNP data sets. Genetics 197, 573–589 (2014).

61. Herrando-Pérez, S., Tobler, R. & Huber, C. D. Smartsnp, an R package for fast multivariate analyses of big genomic data. Methods Ecol. Evol. 12, 2084–2093 (2021).

62. Wang, J. & Zhang, Z. GAPIT Version 3: Boosting power and accuracy for genomic association and prediction. *Genomics*, Proteomics Bioinformatics. 19, 629–640 (2021).

63. Yin, L. et al. rMVP: A memory-efficient, visualization-enhanced, and parallel-accelerated tool for genome-wide association study. *Genomics*, Proteomics Bioinformatics. 19, 619–628 (2021).

64. McLaren, W. et al. The Ensembl Variant Effect Predictor. Genome Biol. 17, 1–14 (2016).

65. Contreras-Moreira, B., et al. Scripting analyses of genomes in Ensembl Plants. In: Edwards, D. (eds) Plant Bioinformatics. Methods in Molecular Biology (ed. Edwards, D.), 2443. 27–55 (New York, NY, 2022). https://doi.org/10.1007/978-1-0716-2067-0_265.

66. Patil, I. Visualizations with statistical details: The ‘ggstatsplot’ approach. J. Open Source Softw. 6, 1–5 (2021).

67. Ksouri, N. et al. Tuning promoter boundaries improves regulatory motif discovery in nonmodel plantsL: the peach example. Plant Physiol. 185, 1242–1258 (2021).

68. Dong, S. S. et al. LDBlockShow: A fast and convenient tool for visualizing linkage disequilibrium and haplotype blocks based on variant call format files. Briefings in Bioinformatics. 1–6 (2020) doi:10.1093/bib/bbaa227.

69. Rangwala, S. H. et al. Accessing NCBI data using the NCBI sequence viewer and genome data viewer (GDV). Genome Res. 31, 159–169 (2021).

