## Supplementary Figures for "ddRAD-seq-derived SNPs reveal novel association signatures for fruit-related traits in peach"

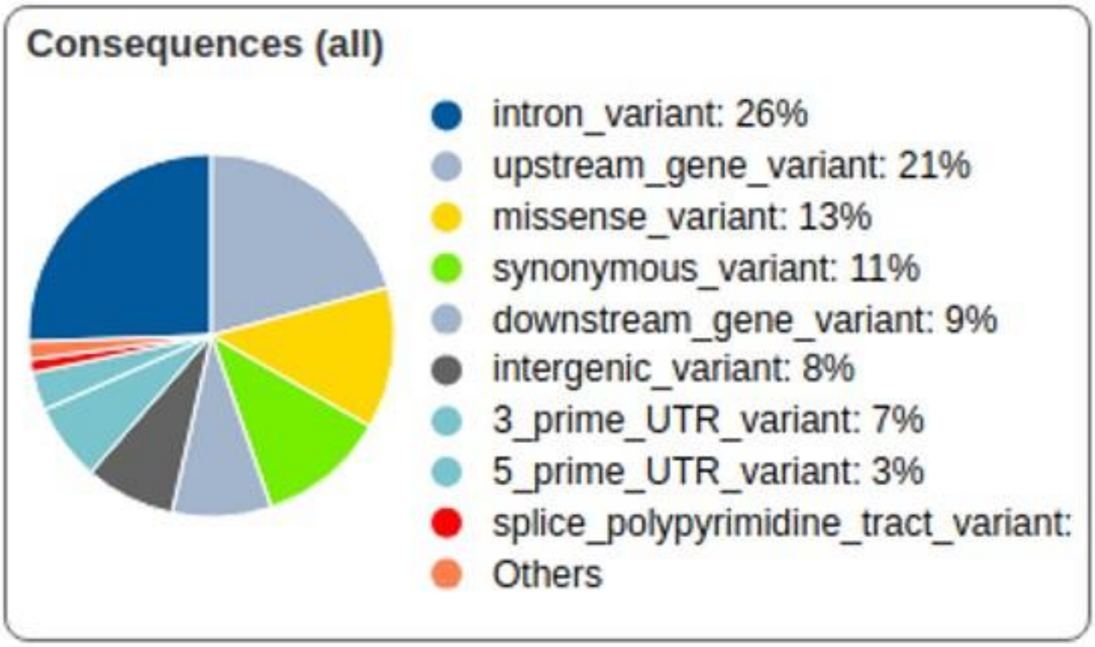


**Figure S1**. Classification of high-quality SNPs based on their genome location. *Prunus persica* genome assembly (GCA_000346465.2) from Ensembl Plant was used as reference. Missense variant corresponds to a change in the codon resulting in a different amino acid while synonymous variant is defined as codon substitution that does not produce a change the encoded amino acid.

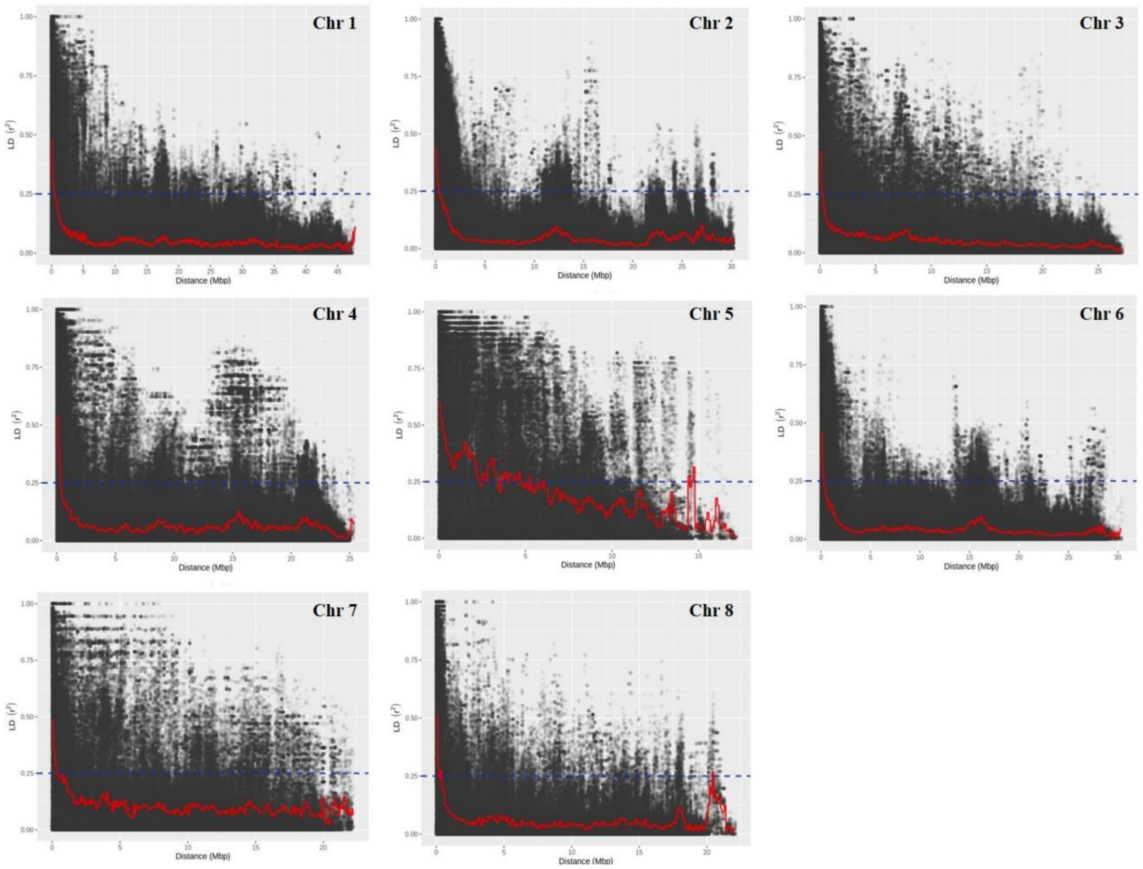


**Figure S2**. Linkage disequilibrium, measured as $r^2$, between pairs of polymorphic marker loci against the physical distance (Mbp). Each dot represents the physical distance between each pair of markers along each chromosome's length.

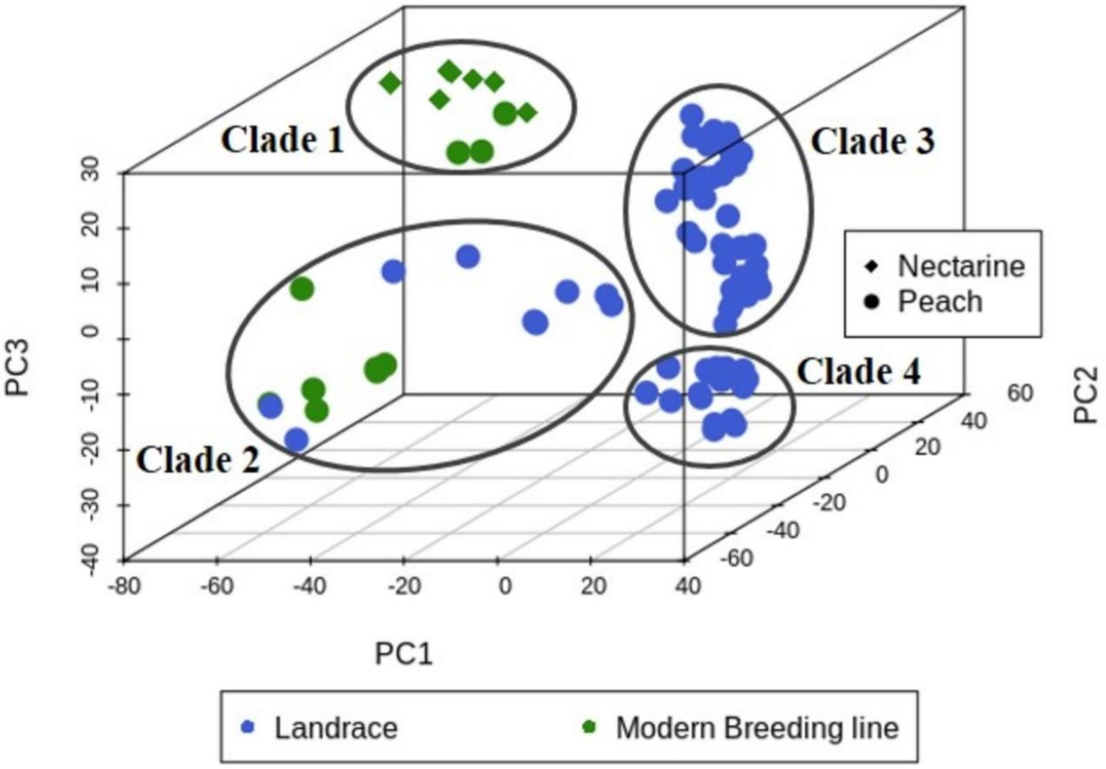


**Figure S3**. Principal component analysis (PCA) of 90 *Prunus persica* accessions. Blue and green colors indicate respectively landraces and modern breeding lines. Shapes correspond to peach and nectarine genotypes. Clades 1 to 4 correspond to the inferred sub-populations (K=4).

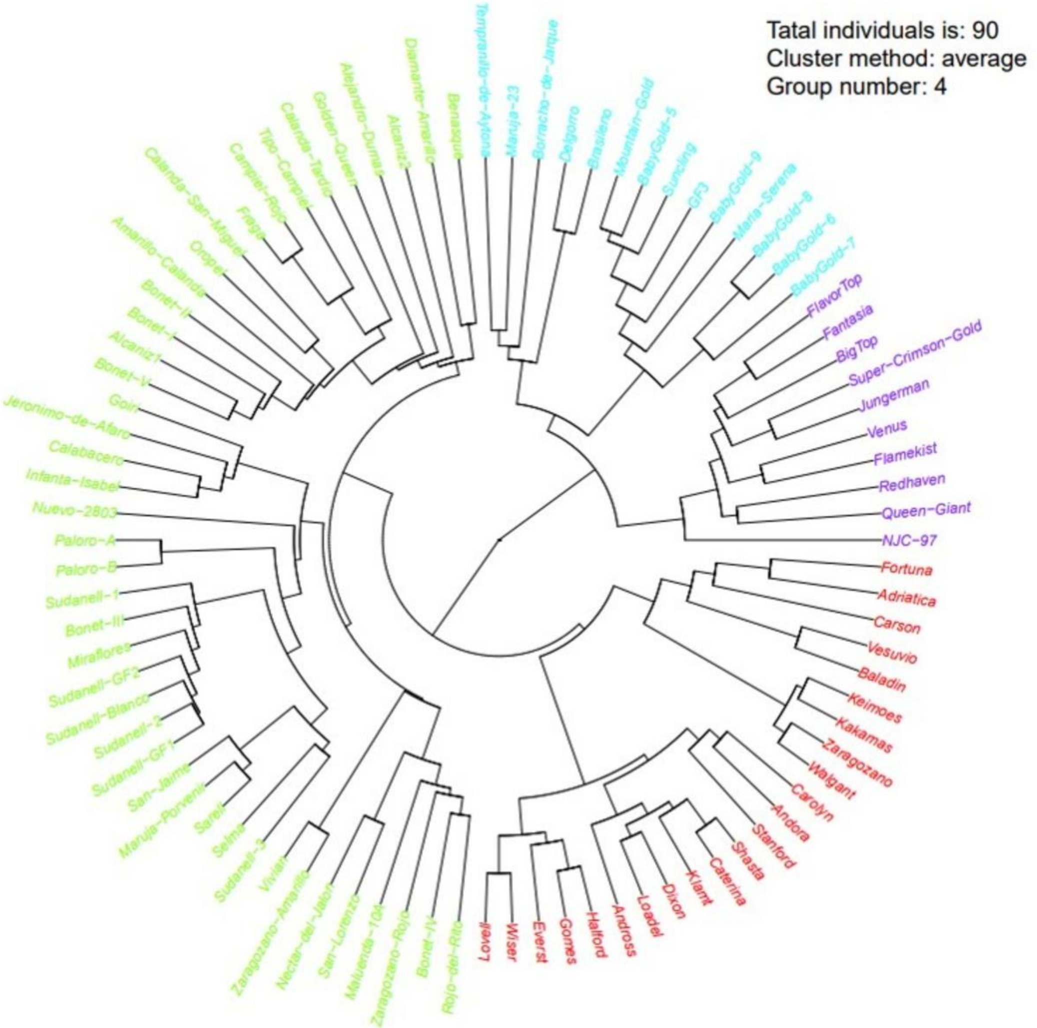


**Figure S4**. The NJ phylogenetic tree of peach and nectarine cultivars. Accession names within each clade are on the outer ring and are depicted in different colors. Purple color corresponds to clade 1 in the PCA plot (Figure S3), blue to clade 2, green to clade 3 and red to clade 4.

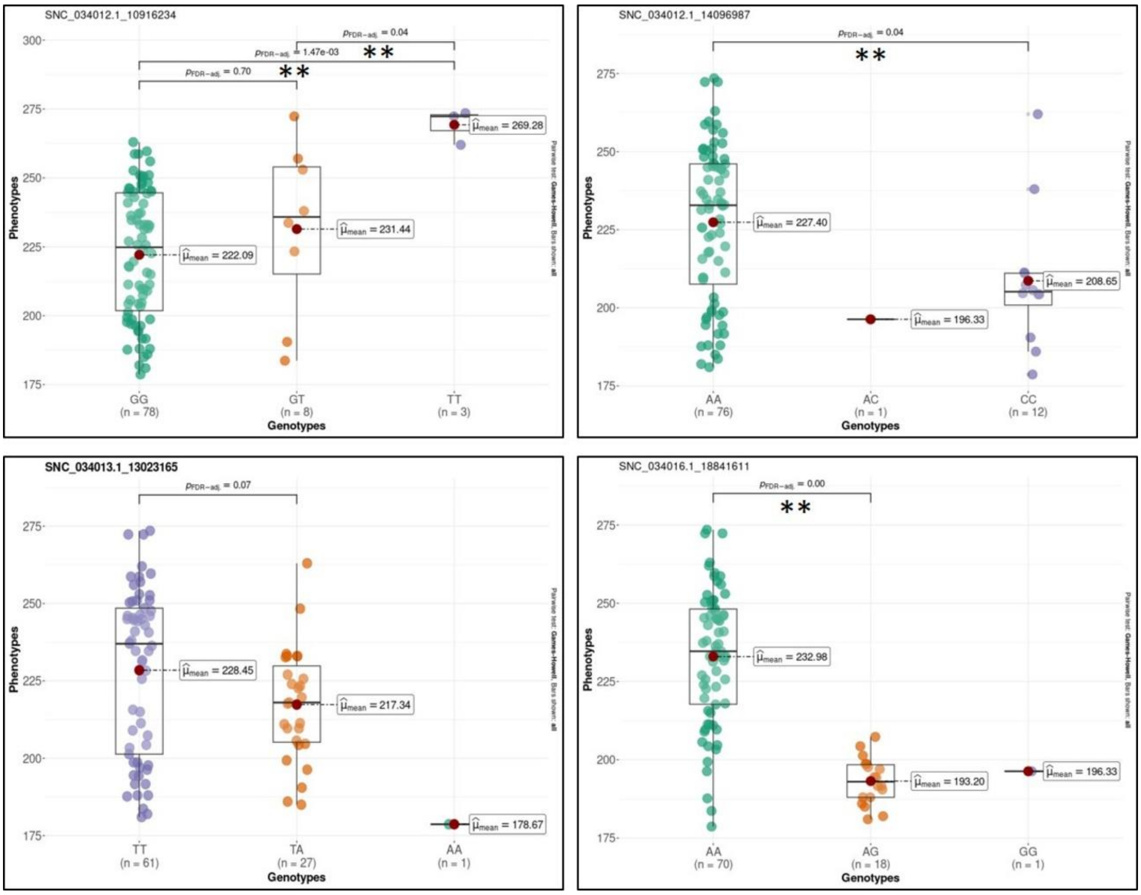


**Figure S5.** Box plot illustrating allelic effect of significant SNPs on harvest date. Y-axis refers to the trait value while x-axis corresponds to the different genotypes (0/0, 0/1 and 1/1). The number of individuals for each genotype is given in parenthesis. Mean values are indicated by red circles and ** indicate significant pairwise comparisons calculated by Games Howel test ($P \leq 0.05$). Lead marker is highlighted in bold.

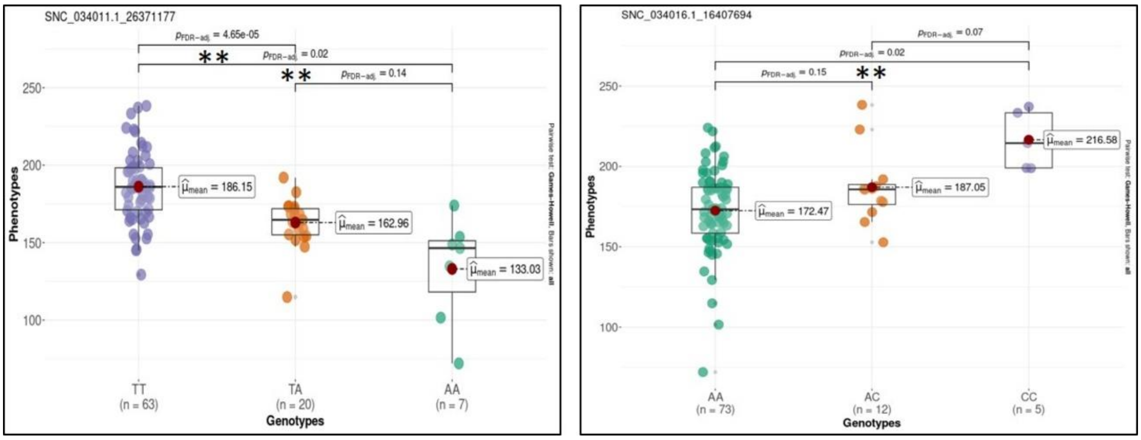


**Figure S6.** Box plot illustrating allelic effect of significant SNPs on fruit weight. Y-axis refers to the trait value while x-axis corresponds to the different genotypes (0/0, 0/1 and 1/1). The number of individuals for each genotype is given in parenthesis. Mean values are indicated by red circles and ** indicate significant pairwise comparisons calculated by Games Howel test ($P \leq 0.05$).

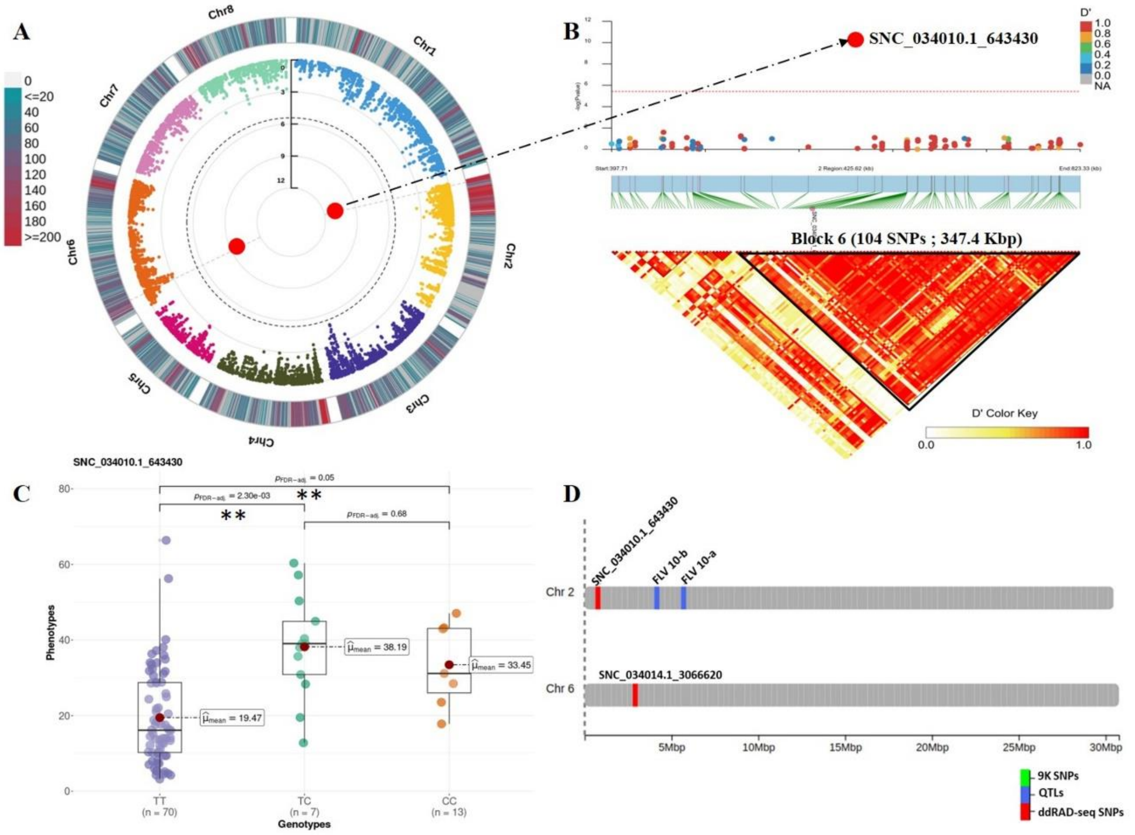


**Figure S7**. Genome Wide Association and LD block analysis for flavonoids content (Flvs). (**A**): Circular Manhattan plot and association signals based on Blink model. Black dashed circular line corresponds to the Bonferroni adjusted threshold (-$\log_{10}(P)$=5.42). Red and large size dots correspond to statistically associated SNPs. Degradation from blue to red indicates the SNP density per 1 Mbp windows across peach chromosomes. (**B**): Locus-specific Manhattan plot (upper panel) and LD heatmap (bottom panel) within 250 kbp on either side of the lead SNP. Pairwise LD measurements are displayed as D' values with a color transition from yellow to red. (**C**): Boxplot depicting allelic effect of lead SNP on trait variation. Mean value for each genotype is indicated by red circle and ** indicate significant pairwise comparisons calculated by Games Howel test ($P \leq 0.05$). (**D**): Genomic distribution of significant ddRAD-derived SNPs (red) and reviewed QTLs in the literature (blue).

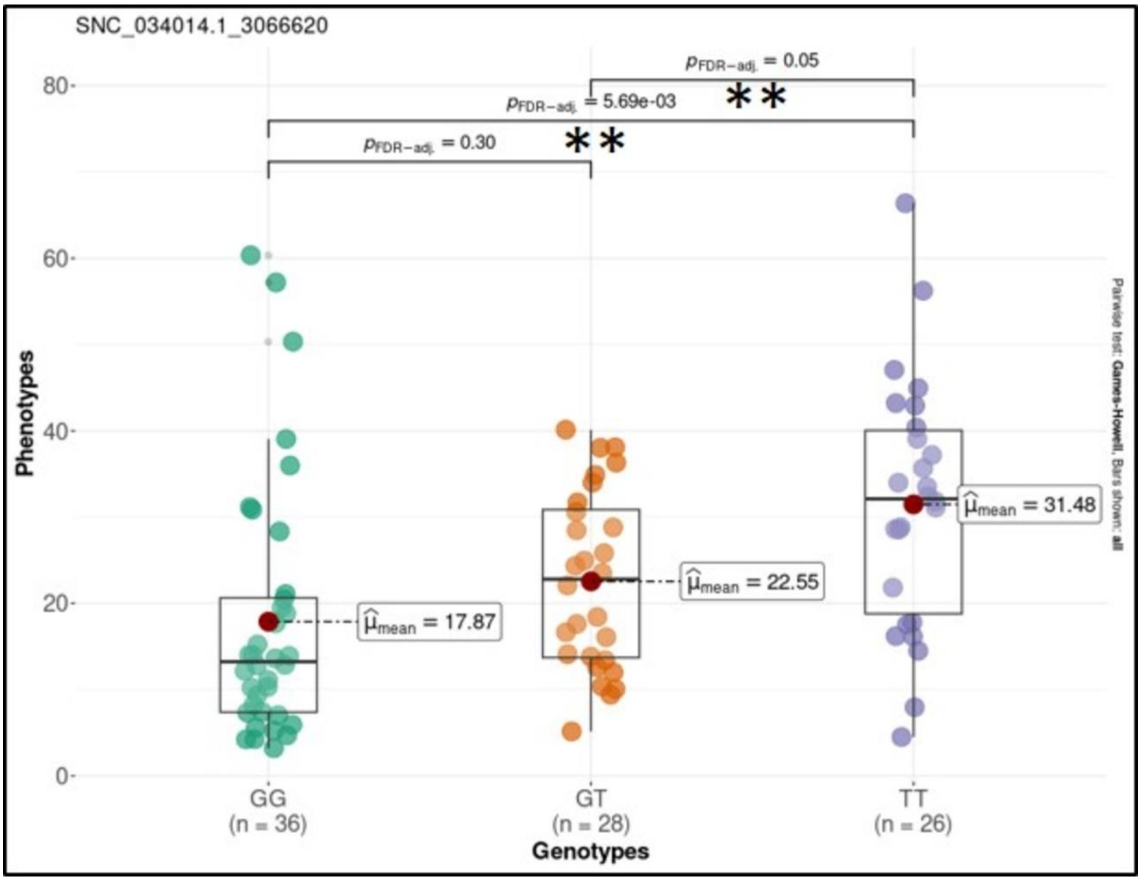


**Figure S8.** Box plot illustrating allelic effect of significant SNPs on flavonoids. Y-axis refers to the trait value while x-axis corresponds to the different genotypes (0/0, 0/1 and 1/1). The number of individuals for each genotype is given in parenthesis. Mean values are indicated by red circles and ** indicate significant pairwise comparisons calculated by Games Howel test ($P \leq 0.05$).

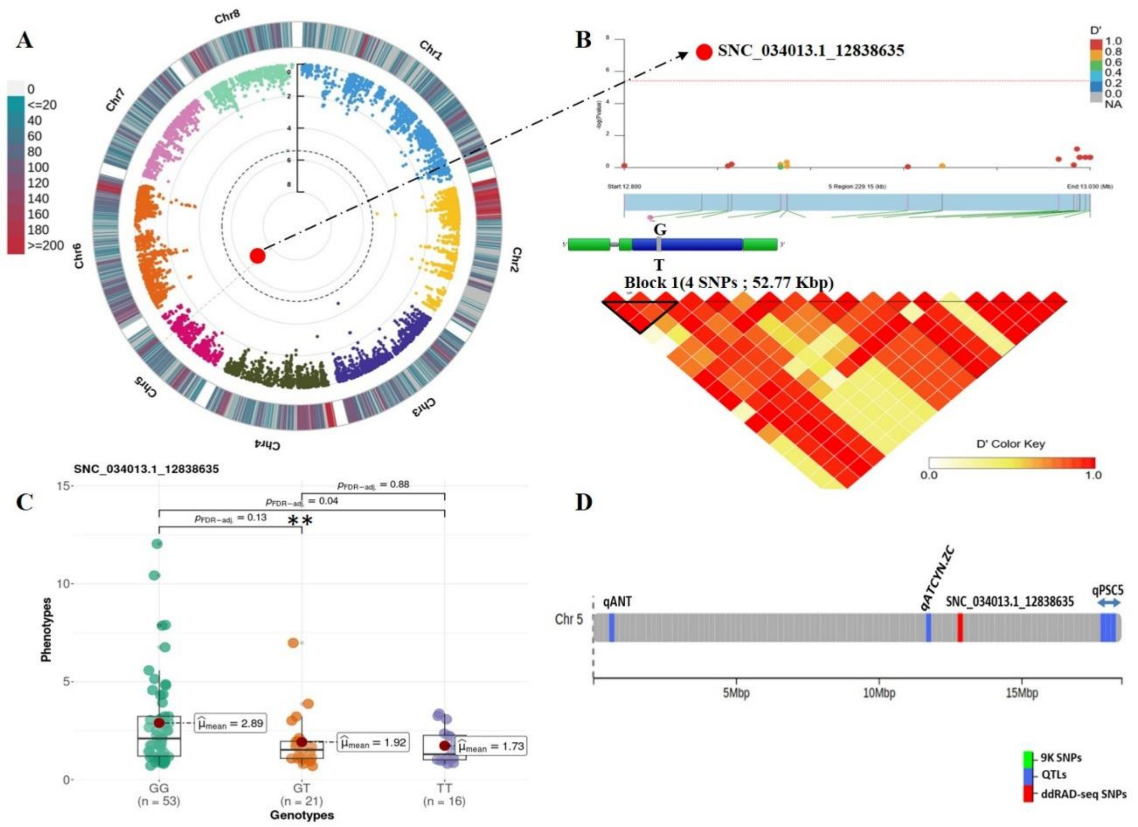

**Figure S9**. Genome Wide Association and LD block analysis for anthocyanin (ACNs). (**A**): Circular Manhattan plot and association signals based on Blink model. Black dashed circular line corresponds to the Bonferroni adjusted threshold ($-log_{10}(P)=5.42$). Red and large size dots correspond to statistically associated SNPs. Degradation from blue to red indicates the SNP density per 1 Mbp window across peach chromosomes. (**B**): Locus-specific Manhattan plot (upper panel) and LD heatmap (bottom panel) within 250 kbp on either side of the lead SNP. Prime candidate gene is represented by blue and green boxes where the blue fragment refers to the exon. Pairwise LD measurements are displayed as D' values with a color transition from yellow to red. (**C**): Boxplot depicting allelic effect of lead SNP on trait variation. Mean value for each genotype is indicated by red circle and ** indicate significant pairwise comparisons calculated by Games Howel test ($P \leq 0.05$). (**D**): Genomic distribution of significant ddRAD-derived SNPs (red) and reviewed QTLs in the literature (blue).

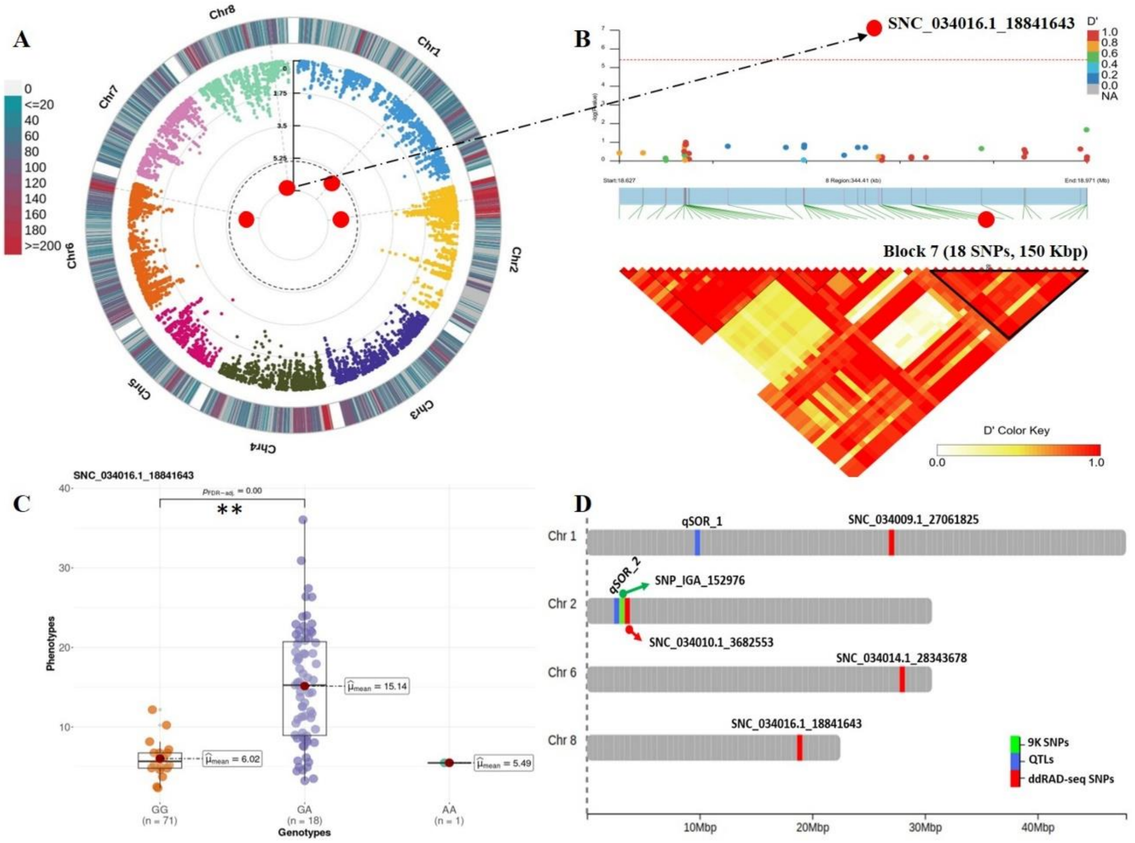


**Figure S10**. Genome Wide Association and LD block analysis for sorbitol content (SRB). (**A**): Circular Manhattan plot and association signals based on Blink model. Black dashed circular line corresponds to the Bonferroni adjusted threshold ($-\log_{10}(P)=5.42$). Red and large size dots correspond to statistically associated SNPs. Degradation from blue to red indicates the SNP density per 1 Mbp window across peach chromosomes. (**B**): Locus-specific Manhattan plot (upper panel) and LD heatmap (bottom panel) within 250 kbp on either side of the lead SNP. Pairwise LD measurements are displayed as D' values with a color transition from yellow to red. (**C**): Boxplot depicting allelic effect of lead SNP on trait variation. Mean value for each genotype is indicated by red circle and ** indicate significant pairwise comparisons calculated by Games Howel test ($P \leq 0.05$). (**D**): Genomic distribution of significant ddRAD-derived SNPs (red), reviewed QTLs in the literature (blue) and 9K array derived SNPs (green).

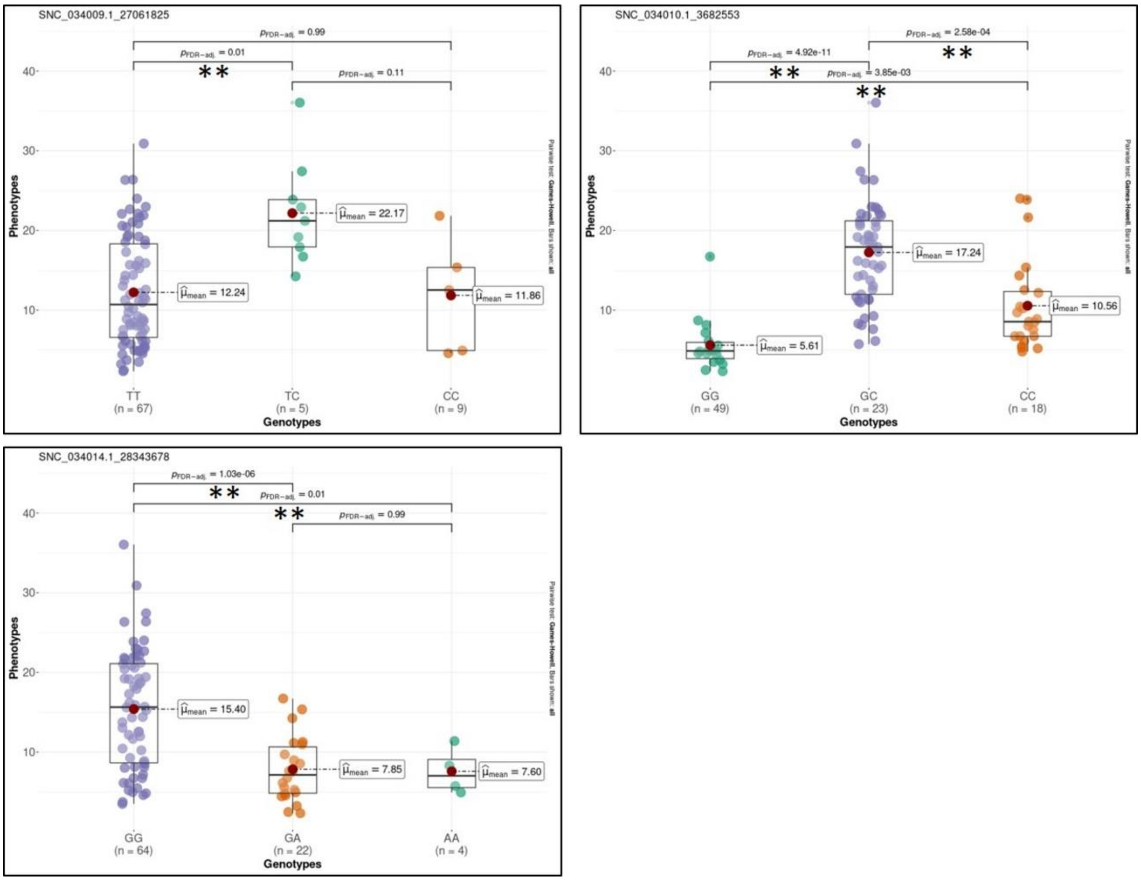


**Figure S11.** Box plot illustrating allelic effect of significant SNPs on sorbitol content. Y-axis refers to the trait value while x-axis corresponds to the different genotypes (0/0, 0/1 and 1/1). The number of individuals for each genotype is given in parenthesis. Mean values are indicated by red circles and ** indicate significant pairwise comparisons calculated by Games Howel test ($P \leq 0.05$).
